# Combining an alarmin HMGN1 peptide with PD-L1 blockade facilitates stem-like CD8^+^ T cell expansion and results in robust antitumor effects

**DOI:** 10.1101/2020.12.15.422990

**Authors:** Chang-Yu Chen, Satoshi Ueha, Yoshiro Ishiwata, Shigeyuki Shichino, Shoji Yokochi, De Yang, Joost J. Oppenheim, Haru Ogiwara, Shungo Deshimaru, Yuzuka Kanno, Tatsuro Ogawa, Shiro Shibayama, Kouji Matsushima

## Abstract

**Background:** The expansion of intratumoral stem-like CD8^+^ T (Tstem) cells provides a potential approach to improving the therapeutic efficacy of immune checkpoint blockade (ICB). Thus, here we demonstrate a strategy to facilitate Tstem cell expansion by combining an alarmin high-mobility group nucleosome binding domain 1 (HMGN1) peptide with programmed death-ligand 1 (PD-L1) blockade.

**Methods:** The antitumor effects of HMGN1, anti-PD-L1 antibody, and their combined treatment were monitored in the B16F10, LLC, Colon26, or the EO771 tumor-bearing mice. The comprehensive immunologic analyses, such as high-dimensional flow cytometry, transcriptome analysis, and single-cell RNA sequencing, were used to investigate the cellular and molecular mechanisms of antitumor immune responses after treatments.

**Results:** The HMGN1 peptide synergizes with PD-L1 blockade in augmenting the number of mature DCs enriched in immunoregulatory molecules (mregDCs) in tumors, and enhancing their MHC class I antigen-presenting program, which is correlated with the expansion of intratumoral Tstem cells, specifically promoting the Tstem cells but restricting terminally exhausted CD8^+^ T (Tex) cells, owing to the regulatory molecules expressed on mregDCs.

**Conclusion:** Our results indicate that HMGN1 peptide serves as an immunoadjuvant to promote effective anti-PD-L1 immunotherapy and implicate that mregDCs play a role beyond facilitating Tstem cell expansion.

## Introduction

Blocking programmed cell death 1 and its ligand programmed death ligand 1 (PD-1/PD-L1) can increase survival in cancer patients by reinvigorating the functions of tumor-reactive CD8^+^ T cells ^1, 2, 3, 4, 5^; however, not all patients respond equally well to PD-1/PD-L1 blockade. The overall response rate to PD-1/PD-L1 blockade alone is below 30% in patients with melanoma, non-small-cell lung carcinoma, ovarian cancer, and renal-cell cancer patients ^6, 7^. Low response rates to ICB therapies are associated with inadequate antigen presentation and insufficient generation of tumor-reactive CD8^+^ T cells ^8, 9^. To address these issues, researchers have attempted to combine various types of immune stimulators with PD-1/PD-L1 blockade to strengthen the antigen-presenting function of intratumoral dendritic cells (DCs) and accelerate the generation of tumor-reactive CD8^+^ T cells, such as R848 (a TLR-7 agonist) ^10^, CpG oligodeoxynucleotide (ODN; a TLR-9 agonist) ^11^ or SD-101 (a novel class of CpG-ODN TLR9 agonist) ^12^. Here, we explore the possibility of combining the immune stimulator called high-mobility nucleosome binding domain 1 (HMGN1) with PD-L1 blockade.

HMGN1 contains two functional domains, a nucleosome binding domain (NBD) and a chromatin unfolding domain (CHUD) ^13, 14^, and functions as both an intracellular and extracellular regulatory protein ^15^. Intracellular HMGN1 is a transcriptional factor that supports higher-order chromatin folding and gene regulation ^13, 14^. Extracellular HMGN1 acts as an alarmin, which is released from apoptotic or necrotic cells, to promote DC activation via MYD88 innate immune signal transduction adaptor (MYD88)- and toll-like receptor adaptor molecule 1 (TRIF)-dependent signaling pathways ^16, 17, 18, 19, 20^. Currently, recombinant HMGN1 is considered to act as a DC adjuvant for T cell-mediated cancer therapy due to its benefits of promoting CD8^+^ T cell expansion ^17, 18, 19, 21^. As reported previously, an *in vitro* assay demonstrated that recombinant HMGN1 can promote DC-dependent CD8^+^ T cell expansion in co-culturing Pmel-1 CD8^+^ T cells with gp100-pulsed BMDCs ^21^. Moreover, an *in vivo* study also demonstrated that HMGN1 synergizes anti-CD4 depleting antibody treatment in enhancing the expansion of intratumoral CD8^+^ T cells in mice ^21^. Thus, we speculate that combining HMGN1 with PD-L1 blockade may facilitate intratumoral CD8^+^ T cell expansion.

Intratumoral CD8^+^ T cell expansion is a promising strategy to improve the therapeutic efficacy of ICB, particularly if it results in the expansion of a newly defined intratumoral CD8^+^ T cell subset with stem-like properties, which acts as resource cells to mediate tumor control in response to ICB therapy ^22, 23, 24^. Stem-like CD8^+^ T (Tstem) cells ^23, 25, 26^, also known as progenitor-exhausted CD8^+^ T (progenitor-Tex) cells ^22, 27^, precursor-exhausted CD8^+^ T cells (T_PEX_) ^24^, and memory-like CD8^+^ T cells ^26, 28^, which are characterized by high expression of SLAM family member 6 (SLAMF6 or also known as Ly108), and transcriptional factor TCF7 (also known as TCF1); the intermediate expression of PD-1; low expression of T-cell immunoglobulin and mucin domain 3 (TIM-3; also known as hepatitis A virus cellular receptor 2, HAVCR2) and lymphocyte-activation gene 3 (LAG-3); and high proliferative capacity ^22, 23, 27, 29, 30^. Patients with melanoma who had increased numbers of Tstem cells displayed durable tumor regression and longer progression-free survival after ICB therapy ^22, 31^. Conversely, a lack of adequate Tstem cells or an abundance of exhausted CD8^+^ T (Tex) cells restricts the therapeutic efficacy of ICB therapy, and limits the survival benefits to both tumor-bearing mice and patients with cancer ^31^.

Here we demonstrate a new strategy to facilitate Tstem cell expansion by combining a HMGN1-derived immunostimulatory peptide with PD-L1 blockade. The HMGN1 peptide synergizes with PD-L1 blockade to increase the number of and the antigen-presenting ability of tumor-infiltrating mregDCs, which is correlated with the expansion of Tstem cells in tumors. Using single-cell RNA sequencing, we predict the cell-cell interaction among mregDC, Tstem cell, and Tex cell populations by CellPhoneDB. The regulatory molecules on mregDCs may support Tstem cell but not Tex cell activation and expansion. Here our findings demonstrate that HMGN1 peptide serves as an immunoadjuvant to promote effective anti-PD-L1 immunotherapy and implicate that mregDCs play a role beyond facilitating Tstem cell expansion.

## Results

### Combined treatment with HMGN1 and PD-L1 blockade induces durable tumor regression

To determine the optimal dosage of murine HMGN1 (mH) in combination with an anti-PD-L1 antibody (αPDL1; clone 10F.9G2), we first performed a dose-response assessment of combined treatment in B16F10 or Colon26 tumor-bearing mice model. After observing the best antitumor effect at 0.08 µg mH in the B16F10 model, and at a dose range of 0.08-0.4 µg mH in the Colon26 model, followed by 200 µg αPDL1 treatment (Fig. 1A, B), we therefore, used the optimized dose of 0.08 µg mH and 200 µg αPDL1 in the following experiments. In this setting, we next evaluated the anti-tumor effects of mH/αPDL1 treatment in Colon26, Lewis lung carcinoma (LLC), EO771, and B16F10 models. Compared with αPDL1 treatment alone, combined treatment of mH/αPDL1 showed significant improvement in tumor growth inhibition in all four different tumor models: Colon26 (on day 17, p < 0.001; day 24, p < 0.01), LLC (on day 18, p < 0.01; day 24, p < 0.01), EO771 (on day 14, p < 0.01) and B16F10 (on day 14, p < 0.05) (Fig. 1A, C).

**Figure 1.**
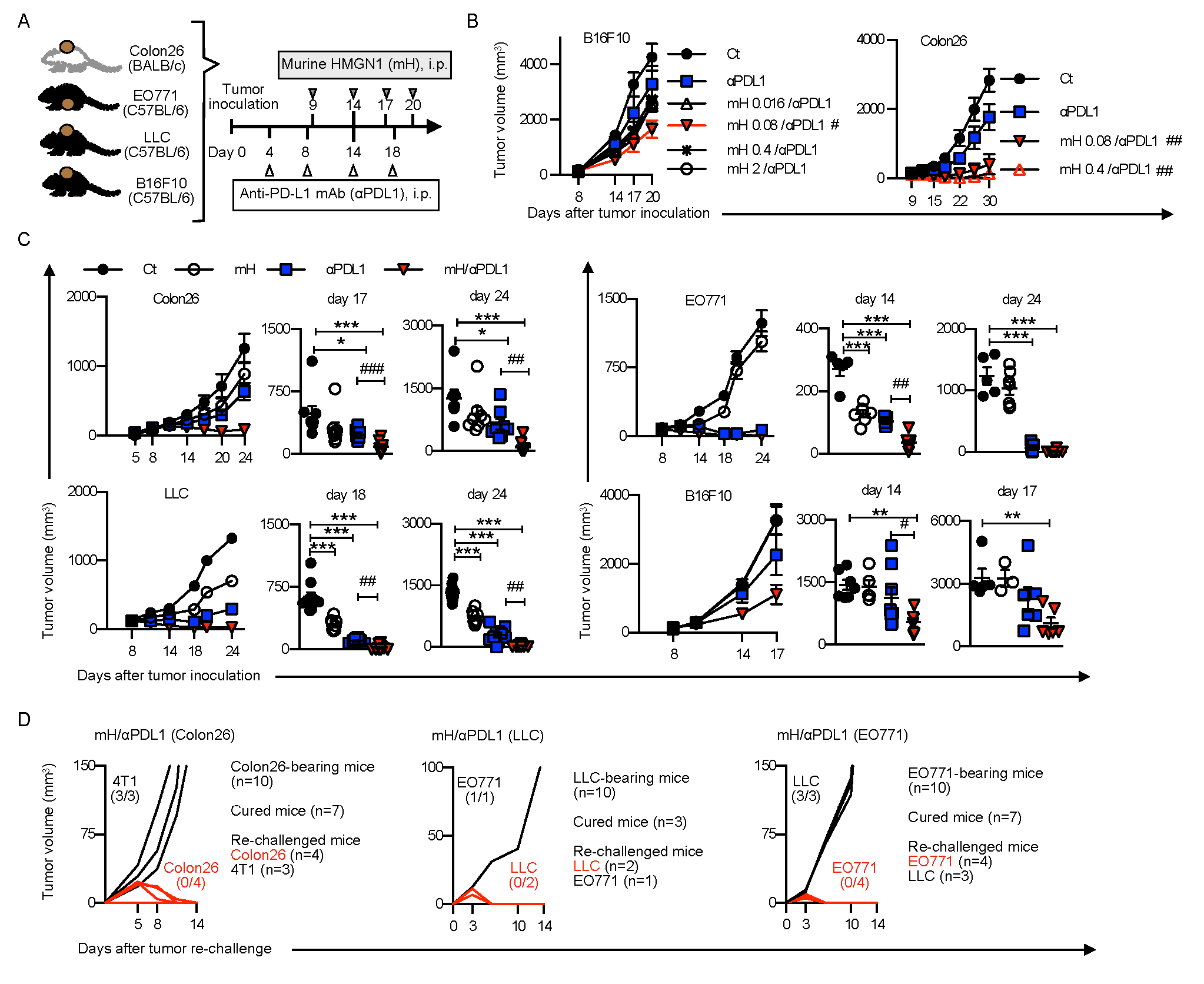
Combined treatment with HMGN1 and PD-L1 blockade induces durable tumor regression. **(A)** The optimized protocol for HMGN1/αPDL1 treatment in Colon26, LLC, EO771, and B16F10 tumor-bearing mice. **(B)** Dose-response for assessing the tumor growth rate of B16F10 or Colon26 tumor-bearing mice treated with combination treatment, in the dose range of 0.016 to 2 μg murine HMGN1. **(C)** Tumor growth analysis during mH/αPDL1 treatment, and the tumor volume of Colon26 (on days 17 and 24), LLC (on days 18 and 24), EO771 (on days 14 and 24), and B16F10 (on days 14 and 17). **(D)** Tumor re-challenge assay. One week after tumor clearance, cured Colon26 tumor-bearing mice were re-challenged with 4T1 and Colon26 tumor cells, cured LLC tumor-bearing mice were re-challenged with EO771 and LLC tumor cells, and cured EO771 tumor-bearing mice were re-challenged with LLC and EO771 tumor cells, respectively. Tumor growth is representative of three independent experiments with at least eight mice per group. Data are presented as mean ± SEM. *, P < 0.05, **, P < 0.01, ***, P < 0.001 for a Dunnett’s post hoc test (compared with control); #, P < 0.05, ##, P < 0.01; ###, P < 0.001 in the figure indicate Student’s *t*-test (comparing between mH/αPDL1-treated and αPDL1-treated groups). Abbreviation: mH (murine HMGN1).

Moreover, we monitored the long-term outcomes and patterns of tumor recurrence after treatments. 70% of Colon26 (seven out of ten), 30% of LLC (three out of ten), and 70% of EO771 (seven out of ten) tumor-bearing mice curing from primary tumors after mH/αPDL1 treatment were also protected from tumor re-challenge (Fig. 1D). These results demonstrated that combining HMGN1 with PD-L1 blockade induces durable tumor regression.

### minP1: a minimized immunostimulatory peptide derived from the HMGN1 NBD retains its anti-tumor effects

As clinical agents, synthetic peptides have advantages over recombinant proteins, as they are quickly and easily synthesized without the risk of endotoxin contamination, and are relatively inexpensive; therefore, we tried to identify the immunostimulatory domain on HMGN1 and used the synthesized HMGN1-derived immunostimulatory domain as a therapeutic peptide in combination with αPDL1 treatment.

In order to determine the immunostimulatory domain on HMGN1, we first synthesized the murine HMGN1 NBD peptide (P1) and CHUD peptide (P2) and evaluated the anti-tumor effects of P1/αPDL1 and P2/αPDL1 treatments in Colon26 model, respectively (Fig.2A). According to the tumor growth assay, P1/αPDL1 treatment, but not P2/αPDL1 treatment, showed synergistic anti-tumor effects equivalent to mH/αPDL1 treatment (Fig. 2B), suggesting that HMGN1 immunostimulatory domain remains within NBD.

**Figure 2.**
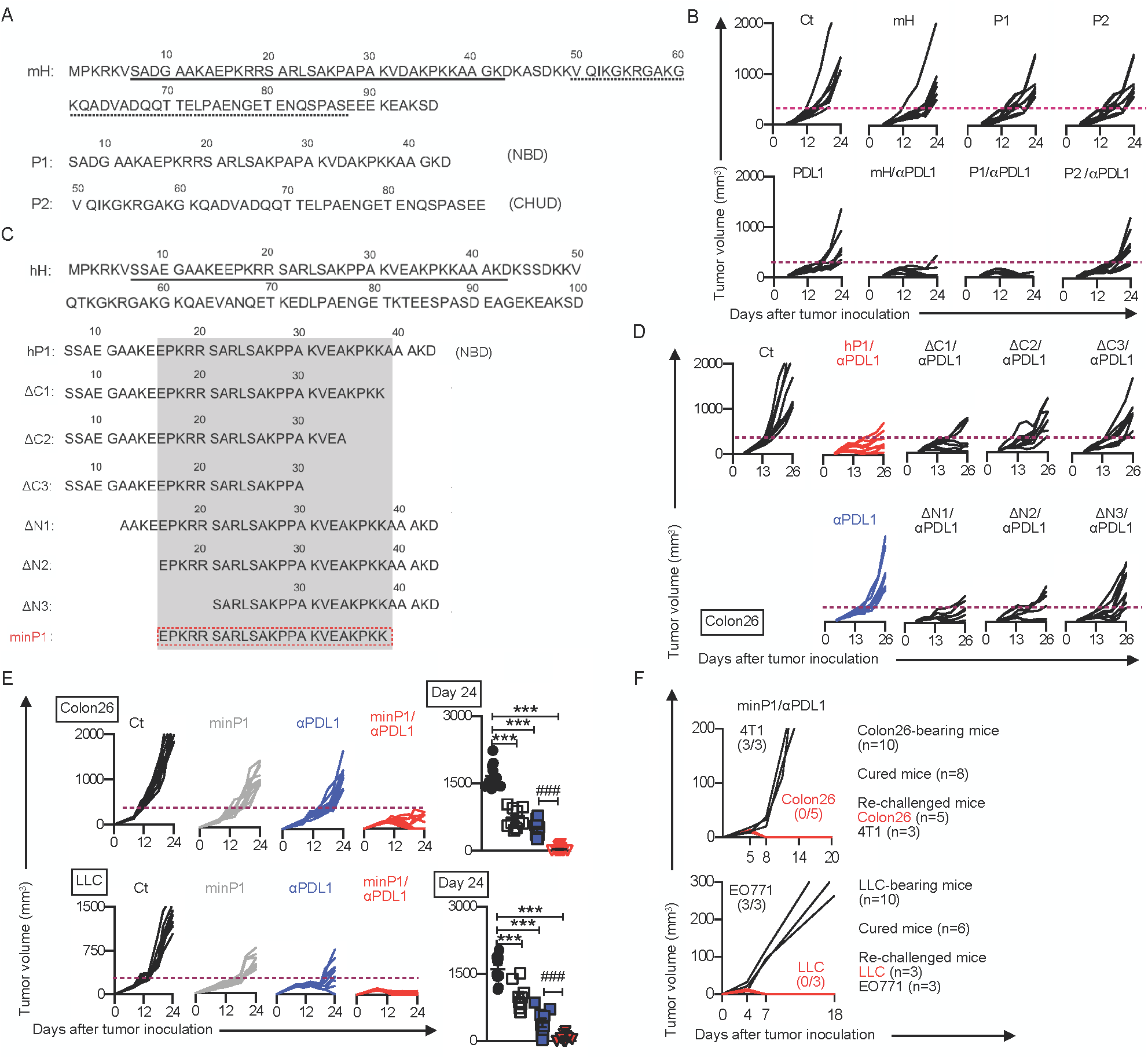
minP1: a minimized immunostimulatory peptide derived from the HMGN1 NBD retains its anti-tumor effects. (**A)** The amino acid sequence of murine HMGN1 (mH) and its nucleosome binding domain (NBD, peptide 1, P1) and chromatin unfolding domain (CHUD, peptide 2, P2). **(B)** The tumor growth analysis on the P1 or P2 in combination with αPD-L1 treatment and the tumor volume in each mouse from day 0 to day 24. **(C)** The amino acid sequence of the human HMGN1 (hH) nucleosome binding domain (NBD) and derived peptides with various lengths. **(D)** Tumor growth analysis after treatment with HMGN1 NBD-derived peptides in combination with αPDL1 treatment and the tumor volume in each mouse. **(E)** Tumor growth analysis after treatment with the minP1(human HMGN1 NBD-derived immunostimulatory peptide) in combination with αPDL1 treatment in both Colon26 and LLC models, and the tumor volume of each mouse from day 0 to day 24. **(F)** Tumor re-challenge assay. One week after tumor clearance, cured Colon26 tumor-bearing mice were re-challenged with 4T1 and Colon26 tumor cells and cured LLC tumor-bearing mice were re-challenged with EO771 and LLC tumor cells, respectively. Tumor growth is representative of three independent experiments with at least eight mice per group. Data are presented as mean ± SEM. *, P < 0.05, **, P < 0.01, ***, P < 0.001 for a Dunnett’s post hoc test (compared with control); #, P < 0.05, ##, P < 0.01; ###, P < 0.001 in the figure indicate Student’s *t*-test (comparing between minP1/αPDL1-treated and αPDL1-treated groups).

Next, in order to minimize the region of immunostimulatory domain within HMGN1 NBD, we further selected human HMGN1 (which has equal anti-tumor efficacy to mH ^21^) and synthesized a human HMGN1 NBD peptide (hP1) and six hP1-derived N-terminally or C-terminally truncated peptides (ΔN1, ΔN2, ΔN3, ΔC1, ΔC2, ΔC3; Fig. 2C). According to the tumor growth assay, the ΔC2, ΔC3, and ΔN3 peptides failed to display synergistic anti-tumor effects in combination with αPDL1 treatment, indicating that the C-terminal region from K_31_ to K_38_ and the N-terminal region from E_16_ to R_20_ are required for function. Thus, we hypothesized that this minimal peptide (minP1, spanning 23-amino acid residues from E_16_ to K_38_, EPKRR SARLS AKPPA KVEAK PKK) retains HMGN1-induced synergistic anti-tumor effects (Fig. 2D).

As expected, minP1 showed equal anti-tumor efficacy as to mH. Combined treatment with minP1 and αPDL1 showed significant improvement in tumor growth inhibition compared with αPDL1 treatment in both Colon26 and LLC models (Fig. 2E). After monitoring the long-term treatment outcomes, we observed 80% of Colon26 (eight out of ten) and 60% of LLC (six out of ten) tumor-bearing mice achieved complete cure from tumors, which showed similar anti-tumor efficacy as to mH. Furthermore, in tumor re-challenging assay, we found that the cured mice resisted primary tumor re-challenge and did not develop secondary tumors (Fig. 2F), consistent with the tumor re-challenge results of mH/αPDL1 treatment (Fig. 1D). Taken together, these data demonstrated that minP1 retains the HMGN1 immunostimulatory function and shows the equal anti-tumor effects to full-length HMGN1 protein.

### minP1 preferentially binds on intratumoral DC populations

In order to further understand how minP1 regulates anti-tumor immunity in tumor-bearing mice, we first tried to identify the primary target population of minP1 therapy in the tumor, secondary lymphoid organ, and primary hematopoietic organ. To address our question, we prepared a fluorescein isothiocyanate (FITC)-conjugated minP1 (FITC-minP1) and took advantage of FITC-minP1 to label the cell populations that would be attached by minP1. 9 days after tumor inoculation, total cell suspensions from the tumor (Tu), draining lymph node (dLN), spleen (SP), and bone marrow (BM) of LLC tumor-bearing mice were pre-stained with an antibody– fluorophore combination and incubated with FITC-minP1. Distinct immune cell populations from Tu, dLN, SP, BM were characterized first (SFig. 1A), and then clustered using t-distributed stochastic neighbor embedding (t-SNE) analysis (SFig. 1B). By visualizing the result of t-SNE analysis, higher FITC-minP1 intensities were detected on monocyte, macrophage, and DC clusters in the Tu, but not detected on any cell clusters in either dLN, SP, or BM (SFig. 1B), which suggests that minP1 may preferentially bind to intratumoral immune cell populations, including monocyte, macrophage, and DC populations.

To confirm the result of t-SNE analysis, we evaluated the FITC-minP1 binding affinity to intratumoral immune cell populations by competitive protein binding assay. Using an unlabeled-minP1 as competitive protein, we found the FITC-minP1 intensities were only decreased on monocyte, macrophage, and DC clusters, but not CD4^+^ T, CD8^+^ T, or NK cell clusters (SFig. 1C). Further in dose-dependent competitive protein binding assay, as the concentration of unlabeled minP1(competitive protein) increased, the amounts of FITC-minP1 bound to macrophage and DC clusters decreased, but the decreasing amounts of FITC-minP1 was not observed on monocyte cluster (SFig. 1D). Moreover, the half-maximal inhibitory concentration (IC_50_) values of DC (69.96 μg/mL) cluster was lower than those for macrophages (192.98 μg/mL), monocyte (>200 μg/mL), and other immune cell populations (>200 μg/mL) in the tumor (SFig. 1D). Taken together, these results demonstrated that minP1 preferentially binds on intratumoral DC populations and may further regulates their biological functions.

### minP1 transcriptionally up-regulates MHC class I antigen presentation program on intratumoral mregDC population

To examine the role in which how minP1 shapes anti-tumor immunity in tumor microenvironment and how minP1 regulates the biological functions of intratumoral DC populations, we analyzed the transcriptomic changes after minP1 treatment by using bulk RNA sequencing (bulk RNA-seq) on whole tumor tissue and single-cell RNA sequencing (scRNA-seq) on CD45^+^ immune cell populations, respectively (Fig. 3A).

**Figure 3.**
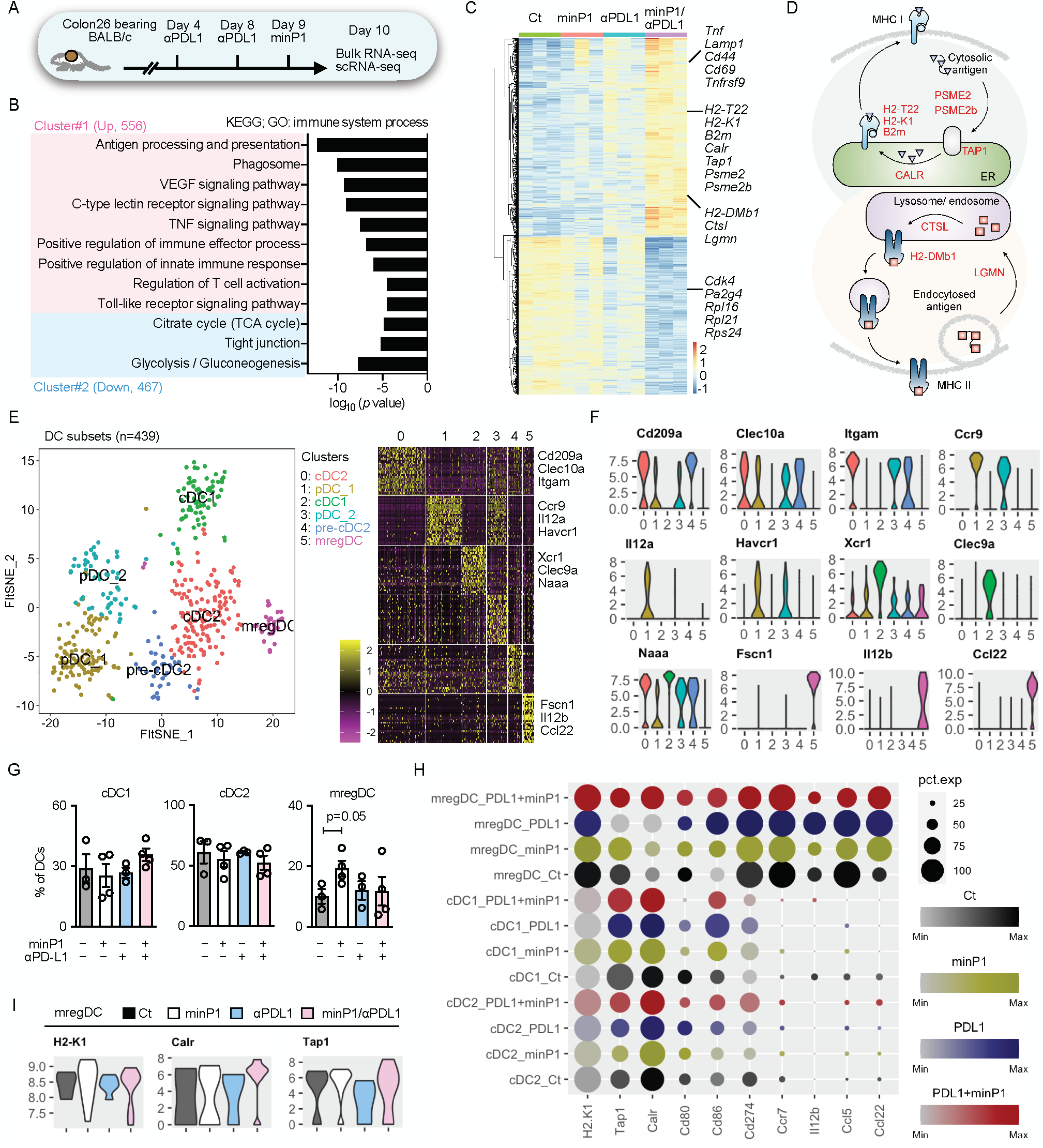
minP1 transcriptionally up-regulates MHC class I antigen presentation program on intratumoral mregDC population. **(A)** Schema on the preparation of tumor cell suspensions from Colon26 tumor-bearing mice for bulk RNA-seq and scRNA-seq on day 10. **(B)** Gene ontology analysis of the differentially expressed genes (DEGs). Each column represents a group, while each row represents an individual gene. The Z-scores of module groups are shown at the bottom-right corner of the heatmap. **(C)** Heatmap of the 1,023 DEGs. **(D)** Summary diagram of antigen processing and presentation pathway components based on the Kyoto Encyclopedia of Genes and Genomes (KEGG) pathway: map04612 (https://www.genome.jp/dbget-bin/www_bget?path:map04612). **(E, F)** A t-distributed stochastic neighbor embedding (t-SNE) projection of scRNA-seq profiling from 439 DCs in tumors of Colon26 tumor-bearing mice. DC clusters are distinct colors. cDC1 (*Xcr1, Clec9a, Naaa*), cDC2 (*Cd209a, Clec10a, Itgam*), pDC (*Ccr9, Il12ra, Havcr1*) and mregDC (*Fscn1, Il12b, Ccl22*) were profiled. Violin plots showing the expression distribution of selected genes in different DC clusters. The y-axis represents log-normalized expression value. **(G)** Proportion of DCs for cDC1, cDC2, and mregDC clusters in different treatment groups. **(H)** Expression of antigen presentation (*H2-K1, Tap1, Calr*) and DC immunostimulatory (*Cd80, Cd86, Cd274*) and cytokine and chemokine (*Ccr7, Il12b, Ccl5, Ccl22*) genes among DC clusters in different treatment groups. Node size is proportional to the expression frequency in a cell cluster. Node color max to min is proportional to the gene enrichment score in each cluster; black represents control (Ct) group. Yellow represents minP1 treatment group. Blue represents PDL1 treatment group. Red represents PDL1+minP1 treatment group. **(I)** Violin plots showing the expression distribution of selected genes (*H2-K1, Calr, Tap1*) in different treatment groups. The y-axis represents log-normalized expression value.

In our bulk RNA-seq data, we found a total of 1,023 differentially expressed genes (DEGs, with adjusted *p* values < 0.05 and a fold-changes of ≥ 2 among groups) identified between αPDL1 and minP1/αPDL1 treatment groups. Gene ontology analysis yielded a large cluster (cluster #1) containing 546 upregulated genes involved in the positive regulation of innate immune responses and immune effector processes, which included genes involved in antigen processing and presentation, phagocytosis, the C-type lectin receptor signaling pathway, the TNF signaling pathway, T cell activation, and the toll-like signaling pathway (Fig. 3B). Interestingly, many upregulated genes in the minP1/αPDL1 treatment group, such as histocompatibility 2, class II, locus Mb1 (*H2-DMb1*), histocompatibility 2, T region locus 22 (*H2-T22*), histocompatibility 2, K1, K region (*H2-K1*), beta-2 microglobulin (*B2m*), transporter 1, ATP-binding cassette, sub-family B (MDR/TAP; *Tap1*), calreticulin (*Calr*), cathepsin L (*Ctsl*), legumain (*Lgmn*), proteasome activator subunit 2 (PA28 beta; *Psme2*), and *Psme2b*, are involved in antigen processing and presentation. Moreover, upregulated genes after combination treatment including tumor necrosis factor (*Tnf*), lysosomal-associated membrane protein 1 (*Lamp1*), *Cd44* molecule, *Cd69* molecule, TNF receptor superfamily, member 9 (*Tnfrsf9*), and IFN gamma receptor 1 (*Ifngr1*), are involved in regulation of T cell activation (Fig. 3C). Conversely, some downregulated genes after combination treatment including cyclin-dependent kinase 4 (*Cdk4*), proliferation-associated 2G4 (*Pa2g4*), and ribosomal proteins 16, 21, and S24 (*Rpl16, Rpl21, Rps24*), are involved in immunosuppression and T cell exclusion ^32^ (Fig. 3C). Our bulk RNA sequencing data demonstrated that minP1 shapes anti-tumor immunity by enhancing antigen presentation and T cell activation programs after αPDL1 treatment in tumor microenvironment.

Furthermore, in our scRNA-seq data, a total of 27 unsupervised cell clusters from tumors were profiled and characterized by unique gene signatures of each cell populations (SFig. 2). In order to inquire deep insight about how minP1 regulates the biological functions on intratumoral DC populations, we selected DC clusters for the following analysis. A total of six DC clusters were profiled and characterized by their unique gene signatures, including conventional DC type1 (cDC1; *Xcr1, Clec9a*, and *Naaa*), conventional DC type 2 and its precursor (cDC2 and pre-cDC2; *Cd209a, Clec10a*, and *Itgam*), plasmacytoid dendritic cells (pDC_1 and pDC_2; *Ccr9, Il12a*, and *Havcr1*), and mregDC (*Fscn1, Il12b*, and *Ccl22*) (Fig. 3E, F). Among those DC clusters, minP1 treatment resulted in a significant increase proportion of mregDC cluster (relative to the untreated group; Fig. 3G), and also enhanced the expression level of MHC class I antigen presentation-related genes: *H2-K1, Calr*, and *Tap1* on mregDC cluster (Fig. 3H, I). Unexpectedly, we did not observe the percentage change or expression change in cDC1 or cDC2 cluster after minP1 treatment (Fig. 3H, I). Taken together, our scRNA-seq data further suggested that minP1 treatment results in an increasing proportion of mregDCs in tumor microenvironment, and also transcriptionally up-regulates MHC class I antigen presentation program on these mregDCs.

### minP1 increases the number of intratumoral mregDCs and enhances their MHC class I expression

To confirm our scRNA-seq result and assess whether the proportion and number of mregDCs increased after minP1 treatments, we prepared total cell suspensions from the tumor tissues of Colon26 tumor-bearing mice on day 12 (after two rounds of minP1 treatments) and 16 (after three times of minP1 treatments) after tumor inoculation. Using flow cytometry, we identified a CCR7^+^ CD11b^+^ CD11c^+^ CD14^−^ Ly6C^−^ MHC class II^+^ cell population as CCR7^+^ mregDC from a lineage-negative cell population (CD3^−^, CD19^−^, Ly6G^−^; Fig. 4A). Increasing numbers of intratumoral CCR7^+^ mregDCs were observed in Colon26 tumor-bearing mice treated with minP1 or minP1/αPDL1 on days 12 and 16, compared to untreated or αPDL1 treatment respectively (*p* < 0.05; Fig. 4B, C). In minP1/αPDL1 treatment group, increased CCR7^+^ mregDCs exhibited higher MHC class I and lower PD-L1 expression compared to those from the untreated or αPDL1 treatment group respectively (*p* < 0.05); however, their MHC class II expression did not exhibit any difference (Fig. 4D), which is consistent with our scRNA-seq results and suggests that minP1 might enhance the MHC class I antigen-presenting ability of intratumoral CCR7^+^ mregDCs. Moreover, consistent with the results in Colon26 tumor-bearing mice, the increasing proportion and the number of intratumoral IL-12b^+^ CCR7^+^ mregDCs ^33^ were also observed in LLC tumor-bearing IL-12b-yellow fluorescent protein (YFP) mice treated with minP1 or minP1/αPDL1, compared to untreated or αPDL1 treatment respectively (*p* < 0.05; Fig. 4E-G).

**Figure 4.**
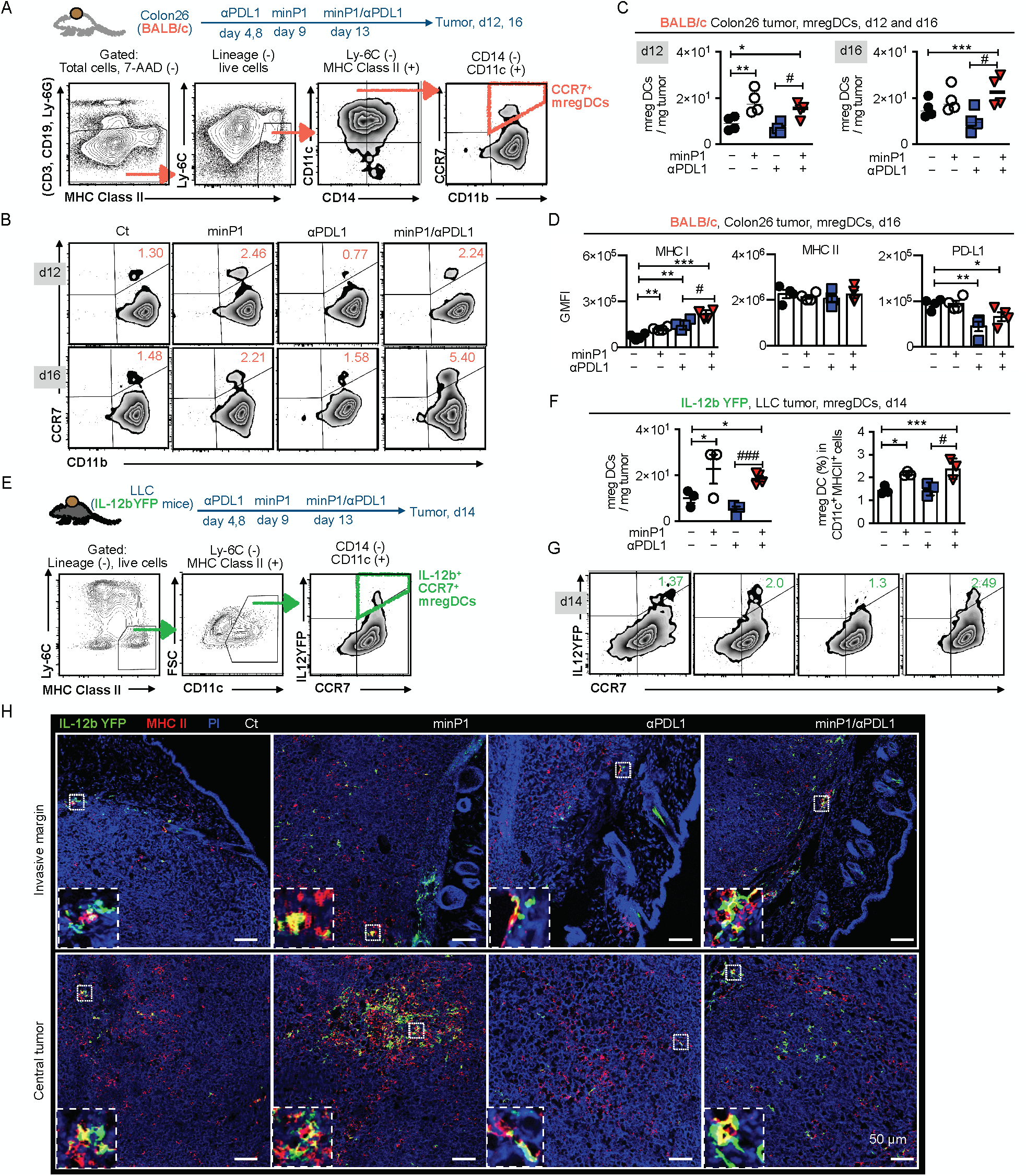
minP1 increases the number of intratumoral mregDCs in both invasive margin and central tumor and enhances their MHC class I expression. **(A)** Schema on the preparation of tumor immune cells from Colon26 tumor-bearing BALB/c mice. Flow cytometry gating of CCR7^+^ mregDCs. **(B, C)** The frequency and number of CCR7^+^ mregDCs in the tumors on days 12 and 16 after tumor inoculation. **(D)** The expression of MHC class I, II, and PD-L1 on CCR7^+^ mregDCs on day16. **(E)** Schema on the preparation of tumor immune cells from LLC tumor-bearing IL12b-YFP reporter C57BL/6 mice. Flow cytometry gating of IL-12b^+^ CCR7^+^ mregDCs. **(F, G)** Frequency and number of the IL-12b^+^ CCR7^+^ mregDCs in the tumor of LLC tumor-bearing IL12b-YFP reporter C57BL/6 mice treated with combination treatment on day 14 after tumor inoculation. **(H)** The number and location of IL-12b^+^ CCR7^+^ mregDCs in LLC tumor sections were identified by co-expression of IL-12b-YFP (green) and MHC class II (red). Each result is representative of three independent experiments with at least four mice per group. Data are presented as mean ± SEM. *, P < 0.05, **, P < 0.01, ***, P < 0.001 for a Dunnett’s post hoc test (compared with control); #, P < 0.05, ##, P < 0.01; ###, P < 0.001 in the figure indicate Student’s *t*-test (comparing with minP1/αPDL1-treated and αPDL1-treated groups). Abbreviation: GMFI (geometric mean fluorescence), 7-AAD (7-Amino-Actinomycin D).

To figure out where these increased intratumoral IL-12b^+^ CCR7^+^ mregDCs locate, we investigated the invasive margins and the central tumors of LLC tumor-bearing IL12-YFP mice. Using immunofluorescence staining, we identified the IL-12b-YFP (green)^+^ and MHC class II (red)^+^ cell population as mregDC in the tumor. Increasing numbers of IL12b^+^ CCR7^+^ mregDCs were observed in both the invasive margins and the central tumors of minP1- and minP1/αPDL1-treated mice (Fig. 4H). Taken together, these results indicated that minP1 enhances the MHC class I antigen-presenting ability of mregDCs and increases the number of mregDCs in both invasive margin and central tumor.

### Expansion of intratumoral stem-like CD8^+^ T cells by minP1/αPD-L1 treatment is associated with the increased number of mregDCs in tumors

The results described earlier with our bulk RNA-seq suggested that minP1/αPDL1 treatment synergistically shapes anti-tumor immunity by promoting regulation of T cell activation program in tumors (Fig. 3B). To confirm our bulk RNA-seq results, we prepared total cell suspensions from the tumor tissues of Colon26 or LLC tumor-bearing mice on days 12 (after twice minP1 treatments) and analyzed intratumoral CD8^+^ T cells in two different type of tumors by flow cytometry (Fig. 5A). The increasing numbers of intratumoral CD8^+^ T cells were observed in minP1/αPDL1 treatment group of both Colon26 and LLC models, compared with αPDL1 treatment group (p < 0.0; Fig. 5B). We next asked whether this synergistic effect on enhancing the expansion of intratumoral CD8^+^ T cells by minP1/αPDL1 treatment was associated with the increased number of mregDCs in tumors. We measured the numbers of mregDCs and CD8^+^ T cells in tumors and found a high correlation between the increasing numbers of mregDCs and CD8^+^ T cells in tumors of both Colon26 (R = 0.78, P = 0.03) and LLC (R = 0.83, P = 0.03) models after minP1/αPDL1 treatment; however, we did not observe this correlation in minP1 (Colon26: R = 0.62, P = 0.13; LLC: R = 0.42, P = 0.39) and αPDL1 (Colon26: R = -0.32, P = 0.69; LLC: R = -0.70, P = 0.11) treatment groups (Fig. 5C). These results suggested that minP1/αPD-L1 treatment brings synergistic effect on enhancing the expansion of intratumoral CD8^+^ T cells, which is associated with the increased number of mregDCs in tumors.

**Figure 5.**
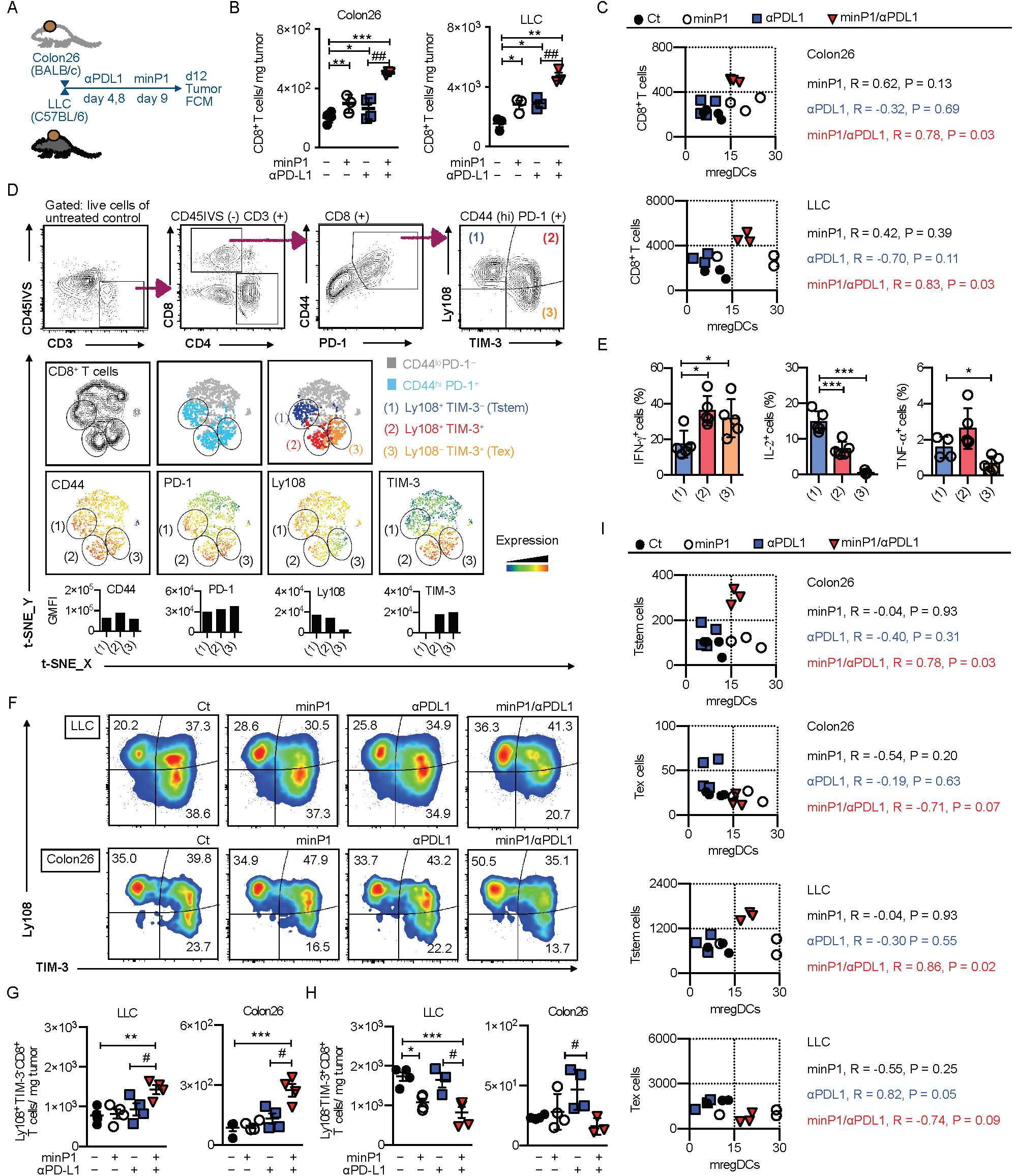
Expansion of intratumoral stem-like CD8^+^ T cells by minP1/αPD-L1 treatment is associated with the increased number of mregDCs in tumors. **(A)** Schema on the preparation of tumor immune cells from Colon26 or LLC tumor-bearing mice. **(B)** The number of CD8^+^ T cells in the tumors of Colon26 or LLC tumor-bearing mice on day 12 after tumor inoculation. **(C)** Pearson correlation coefficients were calculated between numbers of mregDCs and CD8^+^ T cells in tumors. Two-sided P values are shown from a Pearson correlation coefficient r. **(D)** Flow cytometry gating of intratumoral Tstem (Ly108^+^ TIM-3^-^ CD8^+^) and Tex (Ly108^-^ TIM-3^+^ CD8^+^) cells. The tSNE defines the Tstem and Tex populations among CD8^+^ T cells and displays the florescence intensity of CD44, PD-1, Ly108, and TIM-3 in this population. **(E)** Expression of the intracellular cytokines IFN-γ, IL-2, and TNF-α. **(F, G**) The compartments, frequencies, and numbers of Tstem (Ly108^+^ TIM-3^-^ CD8^+^) and Tex (Ly108^-^ TIM-3^+^ CD8^+^) cells in the tumors of LLC or Colon26 tumor-bearing mice on day 12 after tumor inoculation. (**H)** Pearson correlation coefficients were calculated between numbers of mregDCs and either Tstem or Tex cells in tumors. Two-sided P values are shown from a Pearson correlation coefficient r. Each result is representative of three independent experiments with at least four mice per group. Data are presented as mean“± SEM. *, P”< 0.05, **, P < 0.01, ***, P < 0.001 for a Dunnett’s post hoc test (compared with control); #, P < 0.05, ##, P < 0.01; ###, P < 0.001 in the figure indicate Student’s *t*-test (comparing between minP1/αPDL1-treated and αPDL1-treated groups). Abbreviation: CD45IVS: (intravenous CD45 staining)

We next asked whether we could identify which CD8^+^ T cell subsets benefited from this expansion. First, we sought to profile Tstem and Tex cell subsets from intratumoral CD44^high^ PD-1^+^ CD3^+^ CD8^+^ T cells by their expression level of Ly108 and TIM-3 (Fig. 5D). Tstem and Tex cells were characterized as Ly108^+^ TIM-3^−^ and Ly108^−^ TIM-3^+^, respectively. Tstem cells exhibited high IL-2, intermediate TNF-α, and low IFN-γ compared with Tex cells (Fig. 5E), consistent with the previous reports ^22, 23, 27, 29, 30^. In minP1/αPDL1 treatment group, increasing proportion and number of Tstem cells were observed in both Colon26 and LLC models compared to αPDL1 treatment group (p < 0.05, Fig. 5F, G). Conversely, the propotion and number of Tex cells decreased in both models after minP1/αPDL1 treatment (p < 0.05, relative to αPDL1 treatment, Fig. 5F, H). Furthermore, we also checked the correlation between the increasing numbers of Tstem cells and mregDCs in tumors. Surprisingly, there was a high correlation between the increasing number of mregDCs and Tstem in tumors of Colon26 (R = 0.78, P = 0.03) and LLC (R = 0.86, P = 0.02) models after minP1/αPDL1 treatment, but no correlation was observed between the increasing number of mregDCs and Tex (Colon26: R = -0.71, P = 0.07; LLC: R = -0.74, P = 0.09) after minP1/αPDL1 treatment (Fig. 5I). Taken together, these results suggest that an expansion of intratumoral Tstem cells by minP1/αPD-L1 treatment may require the support from the increased number of mregDCs in tumors.

### The multiple regulatory molecules on mregDC that may restrict Tex cell but facilitate Tstem cell activation and expansion

As previously reported, an increasing number of IL12^+^ CCR7^+^ mregDCs correlates with the activation and expansion of intratumoral CD8^+^ T cells, owing to (i) the chemokine receptor CCR7 which enables mregDC to transport antigens from tumor to lymph nodes and then support T cell priming and expansion ^34, 35, 36, 37^, (ii) the cytokine IL-12 which enables mregDC to augment CD8^+^ T cell activation via T cell-DC crosstalk beyond the cytokines IFN-γ and IL-12 ^33^, and (iii) immunoregulatory molecules (PD-L1, PD-L2, CD80, and so on) which enable mregDC control T cell activation ^37^.

However, the mechanisms how mregDCs modulate the activation and expansion of Tstem or Tex cell remain unclear. In order to understand the mechanisms between mregDC (Fig. 3E, F) and Tstem/Tex cells (SFig. 3), we here use CellPhoneDB^38^ to predict their relevant interacting ligand-receptor partners from our scRNA-seq data (Fig. 6A, B). First of all, mregDCs within the tumors express high level of chemokines CCL5, CXCL9, and CXCL16 that may recruit different CD8^+^ T cell subsets into tumor tissue and locally present tumor antigens to re-stimulate those recruited CD8^+^ T cells. The expression pattern of the *CCL5-CCR5* and *CXCL9-CXCR3* ligand-receptor complex between mregDCs and Tstem cells, suggests mregDCs could mediate Tstem cell recruitment. Moreover, mregDCs might also have the ability to recruit Tex cells by other ligand-receptor complexes including *CCL5-CCR5* and *CXCL16-CXCR6* (Fig. 6C, D).

**Figure 6.**
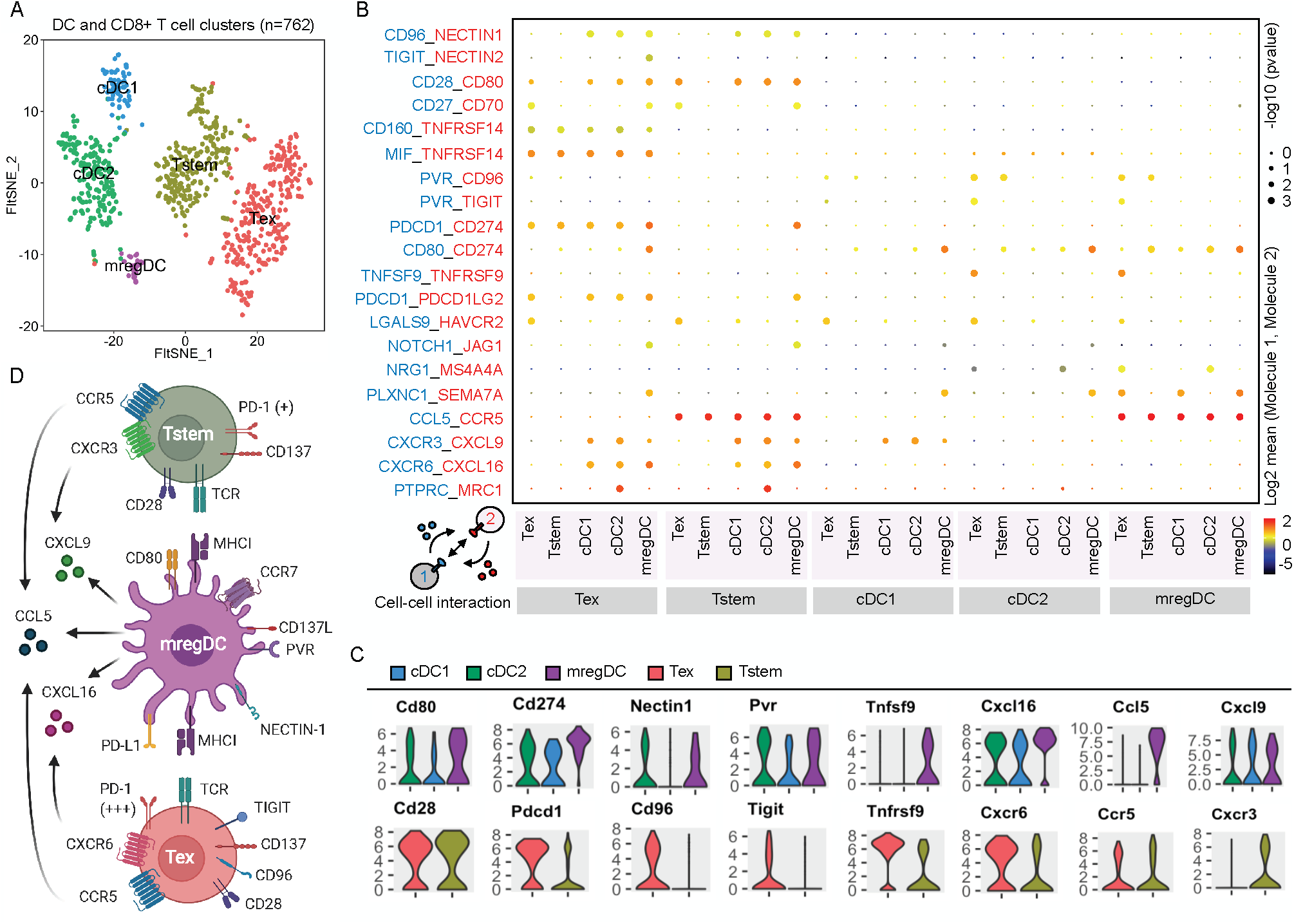
The multiple regulatory molecules on mregDC that may restrict Tex cell but facilitate Tstem cell activation and expansion. **(A)** A t-SNE projection of scRNA-seq profiling from 762 T cells and DCs in tumors of Colon26 tumor-bearing mice. T cell and DC clusters were represented as distinct colors. **(B)** Overview of selected ligand–receptor interactions; P values were indicated by circle size, scale on right (permutation test). The means of the average expression level of interacting molecule 1 in cell cluster 1 and interacting molecule 2 in cell cluster 2 were indicated by color. Assays were carried out at the mRNA level, but were extrapolated to protein interactions. **(C)** Violin plots exhibited the expression distribution of selected genes in T cell and DC clusters. The y-axis represented log-normalized expression value. **(D)** Summary diagram of the main receptors and ligands expressed on the mregDC, Tstem, and Tex cell clusters that were involved in cellular recruitment and co-stimulation/co-inhibition. Created with BioRender.com.

After recruiting Tstem and/or Tex cells, the anti-tumor activity of those T cells might further be regulated by mregDCs within tumors. mregDCs expressed higher immunoregulatory molecules such as *Cd80, Cd274, Nectin1, Pvr*, and *Tnfsf9* (known as CD137 ligand) compared to cDC1 and cDC2 (Fig. 6C). The corresponding receptors or ligands to those immunoregulatory molecules such as *Pdcd1, Cd96*, and *Tigit* were highly expressed on Tex cells but not Tstem cells. By contrast, Tstem cells expressed slightly higher *Cd28* and *Tnfrsf9* (known as CD137) (Fig. 6C). The *CD274-PDCD1, NECTIN1-CD96*, and *PVR-TIGIT* co-inhibition between mregDC and Tex may restrict the activation and expansion of Tex cells (Fig. 6D), conversely, the *CD80-CD28* and *TNFSF9-TNFRSF9* co-stimulation between mregDC and Tstem cells may facilitate the activation and expansion of Tstem (Fig. 6D). Taken together, these results suggest that the multiple regulatory molecules on mregDC that may restrict Tex cell but facilitate Tstem cell activation and expansion.

On account of mregDCs’ possible ability to recruit and regulate both Tstem and Tex cell functions, we further asked whether minP1/αPDL1 treatment could further augment these relevant interacting ligand-receptor partners between mregDC and Tstem/Tex cells which are shown by our CellPhoneDB analysis (Fig. 6B, C). We examined the expression of each immunostimulatory molecules and its corresponding receptor or ligand on mregDC and Tstem/Tex cell populations in different treatment groups. In comparison of untreated control group, *Pvr* and *Cxcl9* expression were slightly upregulated on mregDCs, *Ccr5* expression were upregulated on Tstem cells, and no clearly changed gene expressions could be found on Tex cells in minP1/αPDL1 treatment group (SFig. 4). Those results showed that combination treatment synergistically augmented *CCL5-CCR5, CXCL9-CXCR3*, and *PVR-TIGIT* signaling pathways between mregDC and Tstem/Tex cells, suggests those three signaling pathways may play critical roles in regulating the recruitment and activation of Tstem/Tex cell populations after minP1/αPDL1 treatment.

## Discussion

A newly defined Tstem cells act as a resource cell population mediating tumor control in response to ICB therapy ^22, 31^. It is reported that Tstem cell mainly reside nearby high-dense antigen presenting cell (APC) niches within tumors, and require the support from APCs to maintain their function and differentiation ^39^. For example, adequate number of DCs reinvigorates intratumoral CD8^+^ T cells from exhaustion and support CD8^+^ T cell expansion in tumors by *CD28-CD80* co-stimulation ^29, 40^. In this study, we found that the minP1 synergizes with PD-L1 blockade in increasing the number of mregDCs and enhancing their antigen-presenting function. As recently reported, mregDCs play an immunoregulatory role inside tumors^37^. Our results indicate that mregDCs may restricts Tex but facilitates Tstem cell activation and expansion by multiple immunoregulatory molecules on mregDCs. In particular, Tstem cells with higher levels of *CD28* expression than Tex cells could be more sensitive to *CD28-CD80* co-stimulation ^23^. By contrast, Tex cells with higher levels of *PDCD1, CD96*, and *TIGIT* could be restricted their functions by mregDCs, owing to *PDCD1-CD274, CD96-NECTIN1*, and *TIGIT-PVR* co-inhibition (Fig. 6C, D), which is consistent with the previous reports ^37, 41^. Although we suggested the cell-cell interactions among mregDC, Tstem, and Tex cell populations by analyzing single-cell level, those cell-cell interactions are still unclear and are currently under investigation.

Our work highlights three important implications for both basic and clinical research. First, we have designed an effective strategy for combining the HMGN1 peptide (minP1) with PD-L1 blockade to improve the efficacy of ICB therapy. The use of peptides as therapeutics has advantages of standardized synthesis protocols, low toxicity, and good efficacy. Compared with the HMGN1, minP1 shows a lower risk of endotoxin contamination during synthesis, and could produce in larger quantities within a short time period, and has increasing potential to use as an adjuvant for cancer immunotherapies in the clinic.

Second, the transcriptome signature of tumor tissues after minP1/αPDL1 treatment could be used as an indicator to evaluate clinical response and help determine prognosis of cancer patients. The gene cluster including *H2-DMb1, H2-T22, H2-K1, B2m, Tap1, Calr, Ctsl, Lgmn, Psme2*, and *Psme2b* is involved in antigen processing and presentation, while the cluster including *Tnf, Lamp1, Cd44, Cd69, Tnfrsf9, Ifngr1*, is involved in regulation of T cell activation; and these genes were all upregulated after combination treatment. Conversely, a gene cluster including *Cdk4, Rpl16, Rpl21, Rps24*, and *Pa2g4* was involved in immunosuppression and T cell exclusion, and was down-regulated after combination treatment. These gene clusters could be used as indicators to predict and monitor the efficacy of combination treatment.

Third, we have identified the domain responsible for the immunostimulatory functions of HMGN1. Although previous studies identified the HMGN1 NBD and CHUD based on their distinct intracellular functions ^13, 14^, our previous work has further demonstrated that HMGN1 also has an extracellular function, inducing an immunostimulatory response that results in synergistic antitumor effects ^16, 17, 18, 19, 20, 21^. Nevertheless, the area of the protein mediating the immunostimulatory function was unclear. Here we have found a 23-amino acid peptide within the HMGN1 NBD that retains the immunostimulatory responses and synergistic antitumor effects of HMGN1. This finding will enable in-depth examination of this immunostimulatory domain to understand how HMGN1 binds and interacts with its candidate receptors, such as lymphocyte antigen 96 (LY96; also known as MD2) ^16^ or Gαi protein coupled receptor (GiPCR)^20^.

Overall, our study provides the rationale for combining an HMGN1 immunostimulatory peptide with PD-L1 blockade for cancer immunotherapy.

## Methods

### Mice

Seven-week-old female BALB/c, C57BL/6 were purchased from Japan SLC, Inc. The IL-12-YFP reporter C57BL/6 mice (B6.129-Il12b^tm1.1Lky^/J) were purchased from the Jackson Laboratory (ME, USA). For tumor growth experiment, each group contained 8 mice except where otherwise specified. All animal experiments were conducted in accordance with institutional guidelines with the approval of the Animal Care and Use Committee of The University of Tokyo and Tokyo University of Science.

### Cell lines and tumor models

Murine tumor cell line Colon26 was obtained from the Cell Resource Center for Biomedical Research (RRID: CVCL_0240; Cell Banker, RIKEN BRC, Japan). Lewis lung carcinoma (LLC) was provided from Dr. F. Abe (RRID: CVCL_4358; Nipponkayaku, Tokyo, Japan). B16F10 was obtained from the American Type Culture Collection (ATCC; RRID: CVCL_0159; ATCC, USA). EO771 was obtained from CH3 BioSystems (CVCL_GR23). Colon26 (2 × 10^5^ cells), LLC (2 × 10^5^ cells) and B16F10 (5 × 10^5^) were subcutaneously inoculated into the right flank of mice. EO771 (2 × 10^5^ cells) were implanted into the inguinal mammary fat pad. Tumor diameter was measured by twice weekly and used to calculate tumor volume (V, mm^3^) by using the formula V= L × W × W/ 2 (where L is tumor length and W is tumor width).

### In vivo treatment

Intraperitoneal injections of αPDL1 (200 μg/injection on days 4, 8, 14, and 18; clone 10F.9G2; BioXcell), and HMGN1 and its-derived peptdies (0.08 μg on days 9, 14, 17, and 20) were given to tumor-bearing mice based on the experiment design. Recombinant HMGN1 proteins were produced in *E. coli* and purified using sequential fractionation by heparin affinity column, ion exchange column and reverse-phase column as previously described ^21^. The purity of HMGN1 (>“99%) was confirmed by sodium dodecyl sulfate polyacrylamide gel electrophoresis (SDS-PAGE). The endotoxin level of HMGN1 (< 1 endotoxin unit per mg) was assessed by Endospecy ES-50M Kit (Seikagaku Corporation, Japan). The protein sequence of HMGN1 was identified by Applied Biosystems Procise 492 HT (Thermo Fisher Scientific, USA). The HMGN1 peptides were synthesized by Eurofins Genomics, Japan.

### Flow cytometry

Three minutes before collecting tissues, intravascular leukocytes were stained by intravenous injection of fluorescein isothiocyanate (FITC)-conjugated antibody (3 μg/mouse) against CD45 ^42^. Total cell suspensions were prepared by enzymatic and mechanical dissociation of tissues, as described previously ^43^. Flow-Count Fluorospheres (Beckman Coulter, USA) were used to determine cell numbers. Cells were pretreated with Fc blocking reagent (anti-mouse CD16/CD32 monoclonal antibody, clone 2.4G2; BioXcell), then stained with a mixture of fluorophore-conjugated anti-mouse antibodies. Data were acquired on a CytoFlex flow cytometer (Beckman Coulter, USA) and analyzed by using FlowJo 10.5 software (FlowJo, LLC, USA). Nonviable cells were excluded from the analysis based on forward and side scatter profiles, and dead cells were excluded by propidium iodide (PI) staining. For intracellular cytokine detection, enriched tumor-infiltrating CD8^+^ T cells were re-stimulated with 1 μg/ml ionomycin (IM) and 25 ng/ml phorbol myristate acetate (PMA) in the presence of GolgiPlug (BD Biosciences, USA) for 4 hours at 37 °C. The re-stimulated CD8^+^ T cells were stained with surface antigens, and these cells were stained for intracellular cytokines using a Cytofix/Cytoperm kit (BD Biosciences, USA), according to the manufacturer’s instructions. For the transcriptome analysis, CD8^+^ T cells from the tumor were sorted on FACSAria III Cell Sorter (BD Biosciences, USA).

### Classification of immune cell population by t-SNE

Multicolor flow cytometric data was visualized by T-distributed stochastic neighbor embedding (t-SNE) analysis ^44, 45, 46^, which was used to classify different immune cell populations in various organs in tumor-bearing mice. On 9 day after tumor inoculation, the total cell suspensions from tumor (Tu), draining lymph node (dLN), spleen (SP), and bone marrow (BM) were first stained with a mixture of 12 fluorophores, then incubated with FITC-conjugated minP1 (FITC-minP1). Using FlowJo software, live cells were gated on 7-AAD negative population and minimized to 3×10^4^ cells per sample. The triplicate samples were concatenated into one group and each group were underwent by t-SNE analysis. To identify immune cell populations in the tumor, dLN, and SP, ten parameters (B220, CD3, CD4, CD8, CD11b, CD11c, CD19, I-A/I-E, NK1.1, and Ly6C) were used to separate B cell, CD4^+^ T cell, CD8^+^ T cell, NK cell, NKT cell, monocyte, macrophage/monocyte (monocyte-to-macrophage transition), and DC populations. To identify immune system precursor cells in the BM, eleven parameters (B220, CD11b, CD24, CD115, CD117, CD135, Ly6C, Ly6G, CX3CR1, Sca-1) were used to separate cell populations, including B220^+^ B cell progenitor, Ly6G^+^ granulocyte progenitor, monocyte progenitor (MP), monocyte-dendritic cell progenitor (MDP), common DC progenitor (CDP), Ly6C^+^ monocyte, and Ly6C^−^ monocyte.

### Bulk-RNA sequencing library preparation and analysis

Total RNA was extracted from 30,000 total cells of each tumor tissue, and stored in lysis/binding buffer (Thermo Fisher Scientific, USA). The messenger RNA (mRNA) was isolated from total RNA by Dynabeads M-270 streptavidin with biotin-oligo (dT)25 (Thermo Fisher Scientific, USA). First- and second-strand cDNA synthesis was performed by Superscript reverse transcriptase IV (Thermo Fisher Scientific, USA) and Kapa HiFi DNA polymerase (Roche, Switzerland). While capturing on the beads, cDNA was digested with fragmentase (NEW ENGLAND BioLabs, USA). Each cDNA fragment was ligated to an adapter carrying Ion-Barcode-common sequence 1 (CS1). Finally, an 8-cycle PCR step was performed to enrich for the desired cDNA library molecules, and those cDNA library products were purified by size selection using AMPure XP beads (Beckman Coulter, USA), and were confirmed by Agilent High Sensitivity DNA kit (Agilent Technologies, USA).

The whole transcripts were amplified from total cells of tumor tissue from Colon26-bearing mice and were used to generate the 5’end Serial Analysis of Gene Expression (SAGE)-sequencing libraries. The sequencing was performed by using an Ion 540 Chef kit, an Ion 540 Chip kit, and an Ion S5 Sequencer (Thermo Fisher Scientific) according to the manufacturer’s instructions except the input library concentration was 65 pM. Adapter trimming and quality filtering of sequencing data were performed by using Cutadpat-v2.4 ^47^, Trimommatic-v0.36 ^48^. The filtered reads were mapped on Refseq mm10 using Bowtie2-2.2.5 (parameters: -t -p 11 - N 1 -D 200 -R 20 -L 20 -i S,1,0.50 --nofw). The mapped reads per gene (raw tag counts) were be quantified as gene expression. Between-sample normalization of gene expression was performed against raw count data by using R 3.6.1 (https://cran.r-project.org/) with DESeq2 ^49^ and pheatmap ^50^ packages. Genes with adjusted *p* value less than 0.05 and a fold-change of ≥ 2 between at least three samples were identified as statistically significant differentially expressed genes (DEGs). Raw data from the experiment have been deposited in the NCBI Gene Expression Omnibus (GEO, http://www.ncbi.nlm.nih.gov/geo); accession GSE139291.

Functional analysis of DEGs was performed by using Cytoscape 3.7.1 with ClueGO plugin (v2.5.4) ^51, 52^. Significantly enriched Gene Ontology (GO) terms ^53^ (GO-immune system process, GO levels: 3–8, version: 2019/02/27) and Kyoto Encyclopedia of Genes and Genomes (KEGG, version: 2019/02/27) pathway terms ^54^ in DEGs were explored and grouped, and a term network was constructed based on the overlap of their elements (kappa score = 0.4). Leading terms within each group were defined as the most significantly enriched term in each group. Terms not connected with any other term were excluded.

### Single-cell RNA sequencing library preparation and analysis

For single-cell RNA sequencing (scRNA-seq), we sorted 30,000 CD45^+^ cells from tumors by AutoMACS (Miltenyi Biotec, USA). Sorted cells were stained with TotalSeq mouse Hashtag antibodies (BioLegend, USA). Then, 10,000 labeled cells were trapped and reverse-transcribed using the BD Rhapsody™ (BD, USA) according to the manufacturer’s instructions. For scRNA-seq, cDNA libraries and hashtag libraries were prepared by an optimized process similar to our magnetic bead-based bulk RNA-seq method ^55^. Sequencing libraries were generated by using NEBNext UltraII FS Library Prep Kit (New England BioLabs, USA), and QC of cDNA and final libraries was performed by MultiNA (Shimazu, JP) and qPCR library quantification assay (KAPA). Samples were sequenced on an Illumina Novaseq 6000 S4 flowcell (67 bp read 1 and 140 bp read 2) (Illumina, USA) to a depth of approximately 100,000 reads per cell. For mouse data, after adaptor removal by using Cutadpat-v2.10 ^47^, gene-expression libraries were aligned to the mm10 reference transcriptome by Bowtie2-2.3.4.1, and count matrices were generated using the home-built shell scripts and the modified python script of BD Rhapsody workflow. Valid cell barcodes were identified as cell barcodes above inflection threshold of knee-plot of total read counts of each cell barcode identified by DropletUtils package ^56^. Each sample origin and doublets were identified based on fold-change of the normalized read counts of Hashtag antibodies. Data were log-transformed [log(TPM + 1)] for all downstream analyses, most of which were performed using the R software package Seurat v2.3.4 ^57^ (http://satijalab.org/seurat) to analyze immune cell populations.

For each cell, three quality control metrics were calculated: (1) the total detected number of genes; (2) the proportion of ribosome encoded transcripts; and (3) the proportion of mitochondrially encoded transcripts. Cells were excluded from downstream analysis if fewer than 200 genes were detected. There was an expression matrix of 7,613 cells by 22,833 genes. Each gene expression measurement was normalized by total expression within the corresponding cell and multiplied by a scaling factor of 1,000,000. Highly-variable genes identified by Seurat package were used for principal components analysis. Principal components were determined to be significant (P < 0.05) using the jackstraw method and Fit-SNE 58 was performed on these key principal components. Unsupervised clustering was performed using a shared nearest neighbor modularity optimization-based algorithm, as described previously ^59^. In our initial analysis of DC and T cell subsets (SFigure. 2), we observed contaminating subpopulations and removed them from the dataset by proceeding with the re-clustering analysis. Differential gene expression and signature enrichment analyses were performed by a Wilcoxon rank-sum test. To perform an analysis of cell-cell interaction among DC and T cell subsets, we analyzed our scRNA-seq data by CellPhoneDB ^38^ (www.CellPhoneDB.org). As previously reported ^60^, CellPhoneDB could calculate the means of the average expression level of each receptor–ligand pair in each pairwise comparison between two cell types, and provide a *P* value for each receptor–ligand pair. We then determined the order of interactions that are highly enriched between cell types based on the number of significant pairs, and manually selected biologically relevant ones. The means of the average expression level of interacting molecule 1 in cell cluster 1 and interacting molecule 2 in cell cluster 2 were indicated by color and *P* values were indicated by circle size, scale. Assays were carried out at the mRNA level, but are extrapolated to protein interactions.

### Immunofluorescent staining

Acetone-fixed, 8-μm tumor sections of LLC tumor-bearing IL12-YFP reporter C57BL/6 mice on day 14 after tumor inoculation, were prestained with anti-mouse MHC class II (clone M5/114.15.2, BioLegend) and incubated with PI. Sections were mounted with ProLong Gold Antifade Mountant (Thermo Fisher Scientific, USA) and examined under a TCS SP5 confocal microscope (Leica Microsystems, Wetzlar, Germany).

### Quantification and statistical analysis

Data were analyzed using Prism 8.0 software (GraphPad Software, USA). For comparisons among groups in the *in vivo* study, we used one-way ANOVA with the Dunnett’s test. For comparisons between the means of two variables, we used two-sided unpaired Student’s *t*-test. For correlation, we used a two-sided Pearson correlation with coefficient r. All statistical analyses were presented as mean with SEM, and conducted with a significance level of α = 0.05 (P< 0.05).

## Acknowledgements

The authors would like to thank Shin Aoki, Shunichi Fujita, and Ai Yamashita (The University of Tokyo, Tokyo) for help with maintenance of laboratory and animal facility, Rina Matsukiyo and Kazushige Shiraishi (Tokyo University of Science, Chiba) for help with cell sorting, Junko Yasuda (Tokyo University of Science, Chiba) for help with RNA sequencing and analysis. C.Y.C. was supported by the University of Tokyo Fellowship. Research work in the laboratory of K.M. is funded by Grant-in-Aid for Scientific Research on Innovative Areas (Grant Number: 17929397).

## Author contributions

Conceptualization, C.Y.Chen, S.Ueha, J.J.Openheim, S.Shibayama, and K.Matsushima; Methodology, C.Y.Chen, S.Ueha, S. Shichino, Y.Ishiwata, S.Yokochi, H.Ogiwara, S.Deshimaru, Y. Kanno, and T. Ogawa; Investigation, C.Y.Chen, S.Ueha, S. Shichino, Y.Ishiwata, De Yang, H.Ogiwara, S.Deshimaru, and K.Matsushima; Writing – Original Draft, C.Y.Chen; Review & Editing, C.Y.Chen, S.Ueha, J.J.Openheim, and K.Matsushima; Funding Acquisition, S.Ueha. S.Shibayama, and K.Matsushima; Supervision, S.Ueha, and K.Matsushima.

## Reporting summary

Further information on research design is available in the Nature Research Reporting Summary liked to this article.

## Availability of data and materials

The datasets used and/or analyzed during the current study are available from the corresponding author on reasonable request.

## Research Reporting Summary

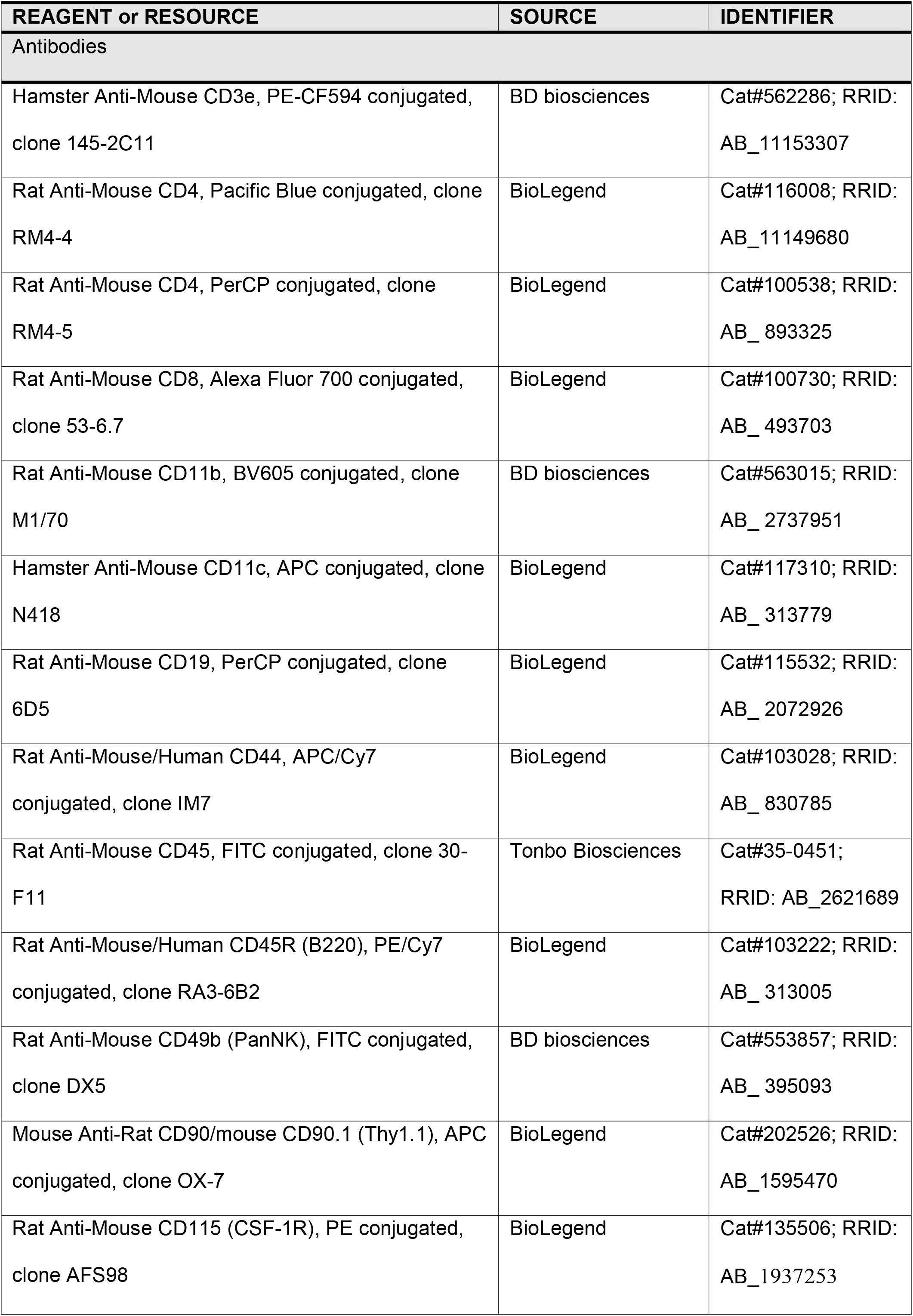
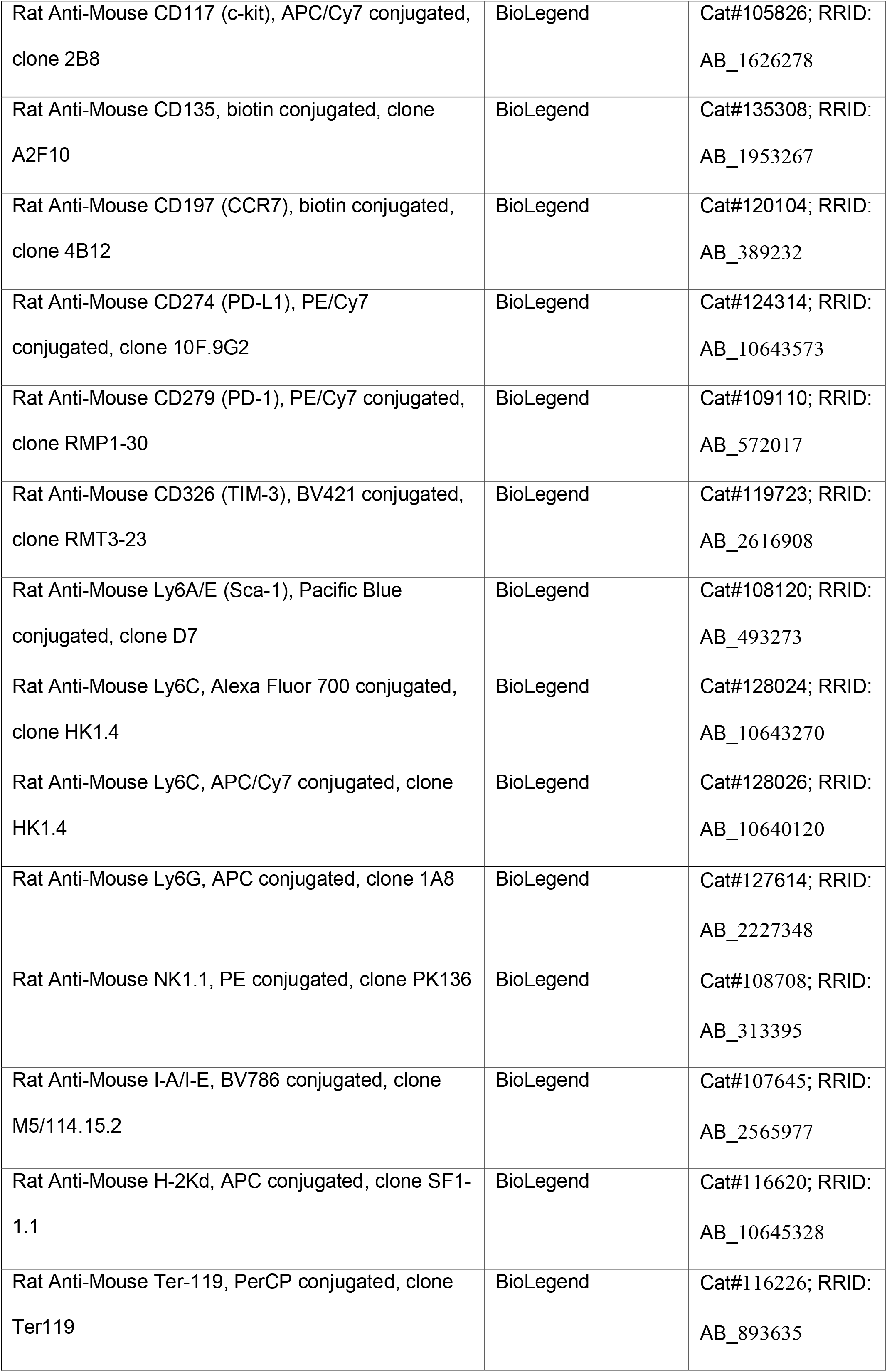
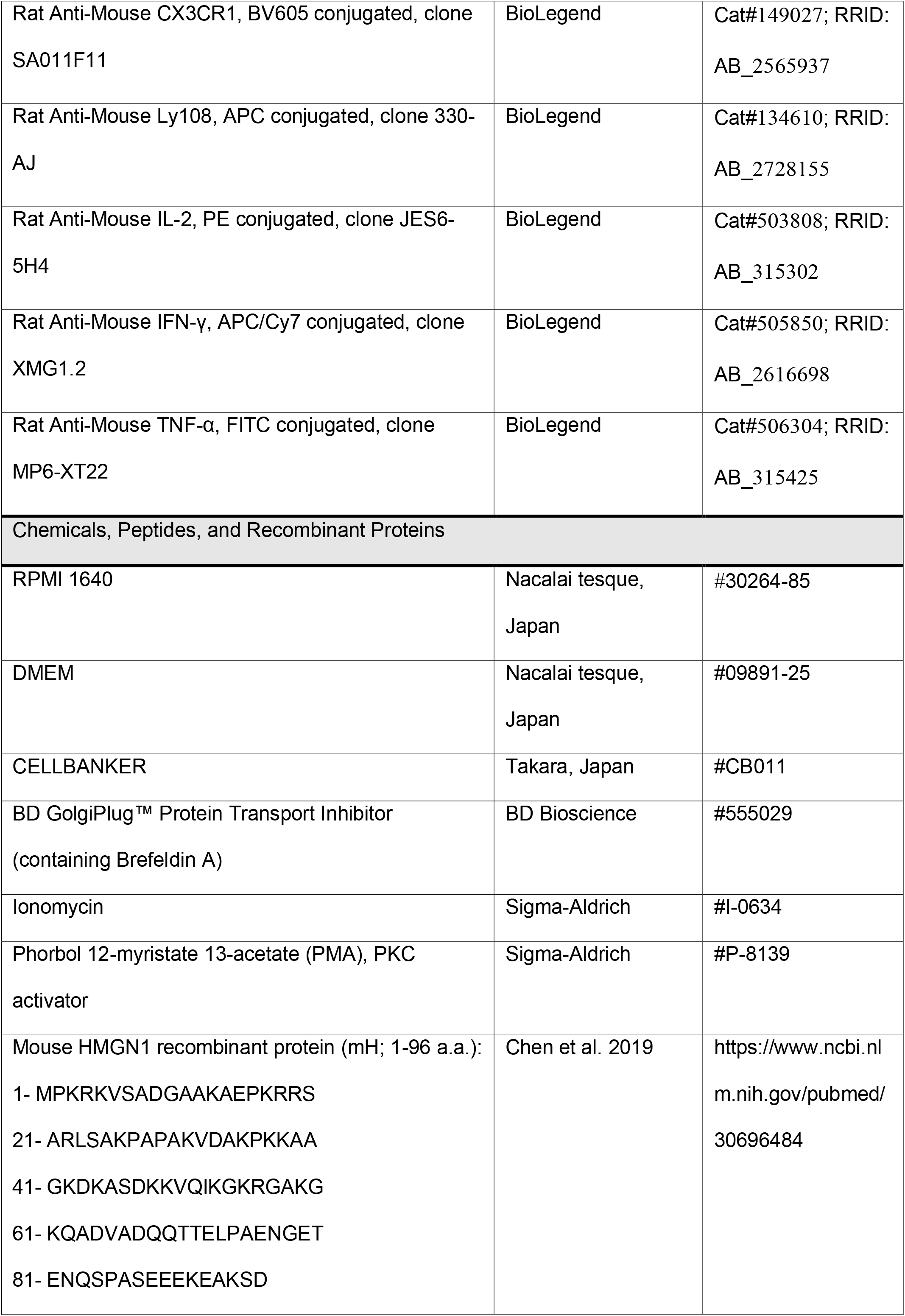
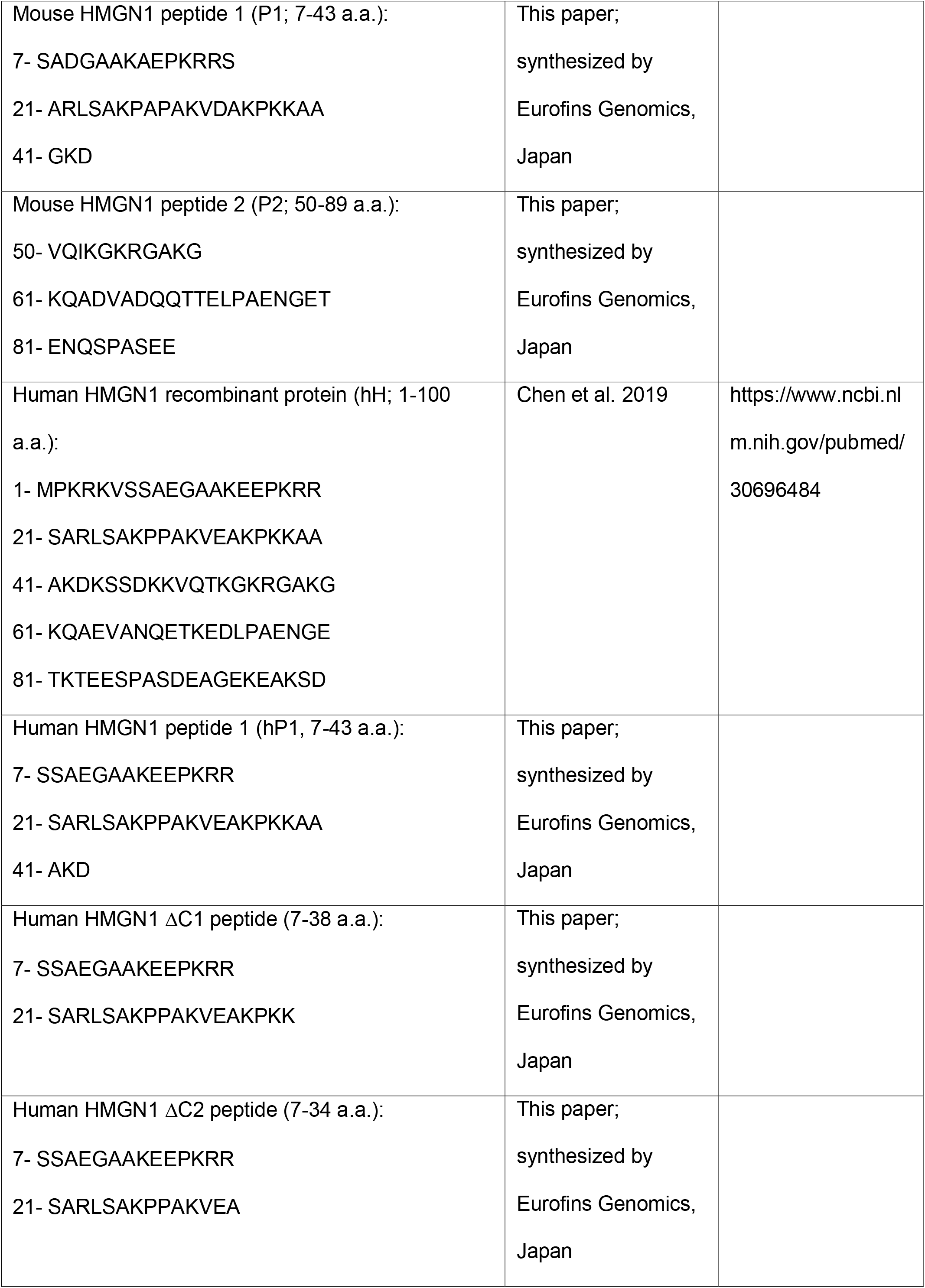
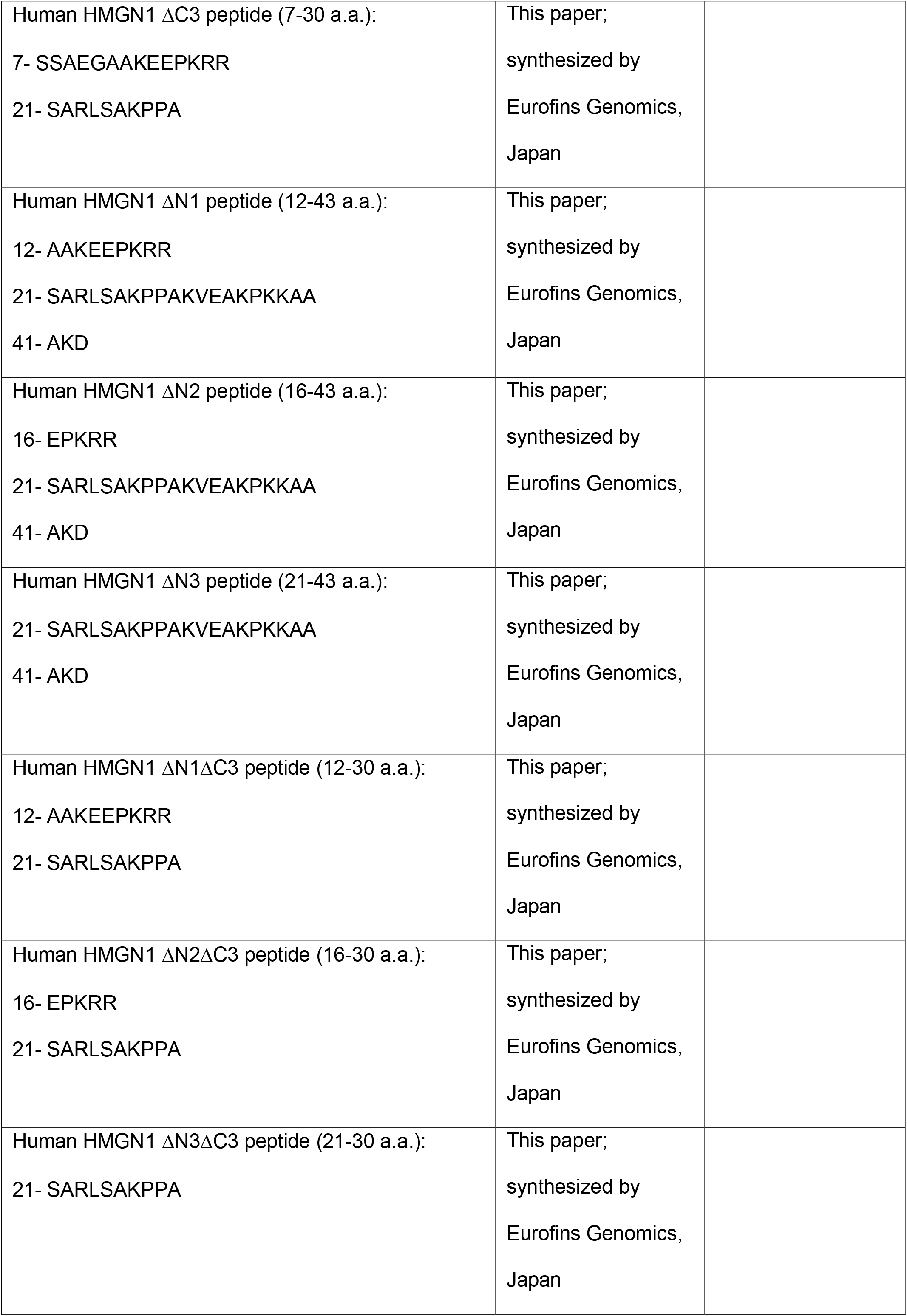
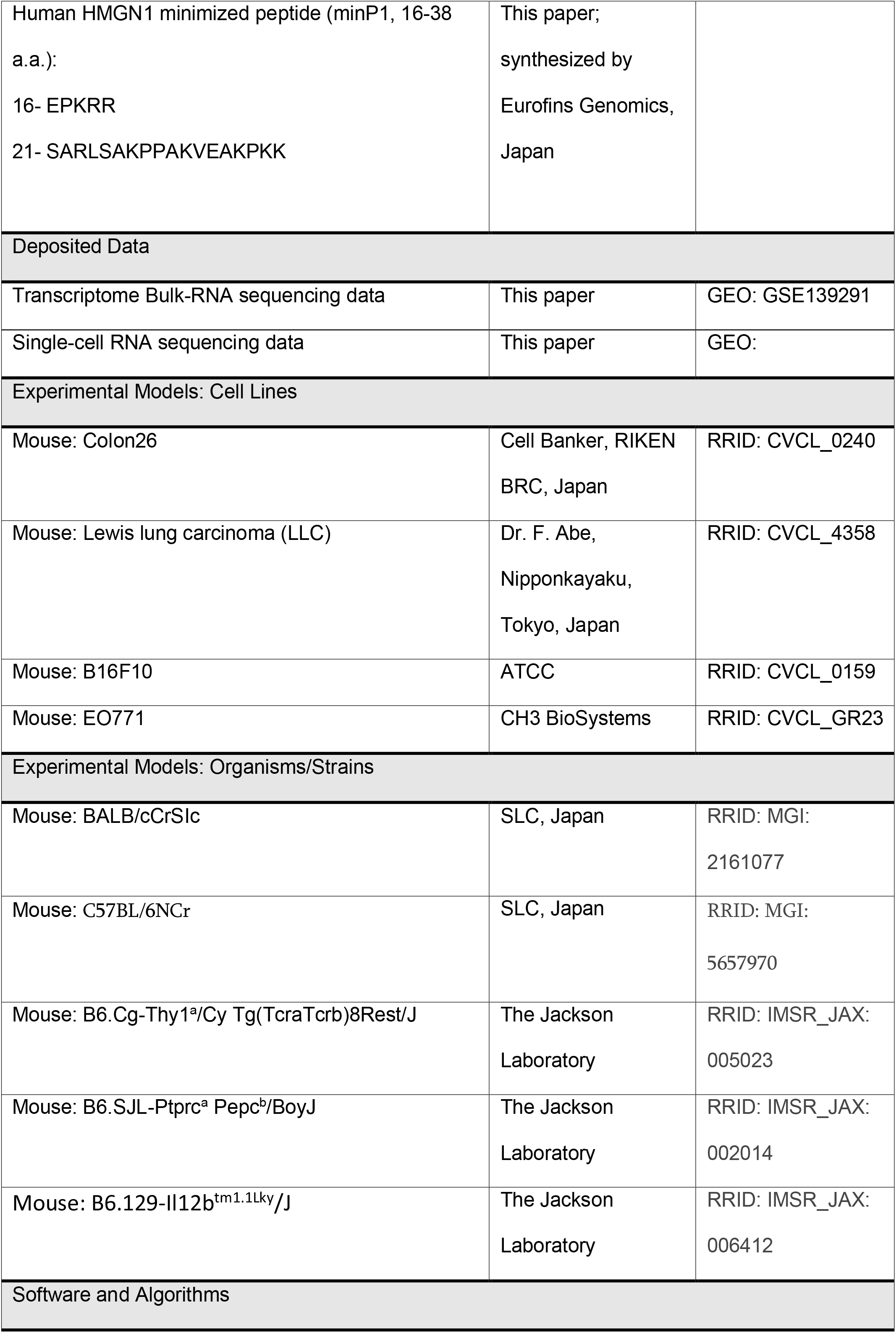
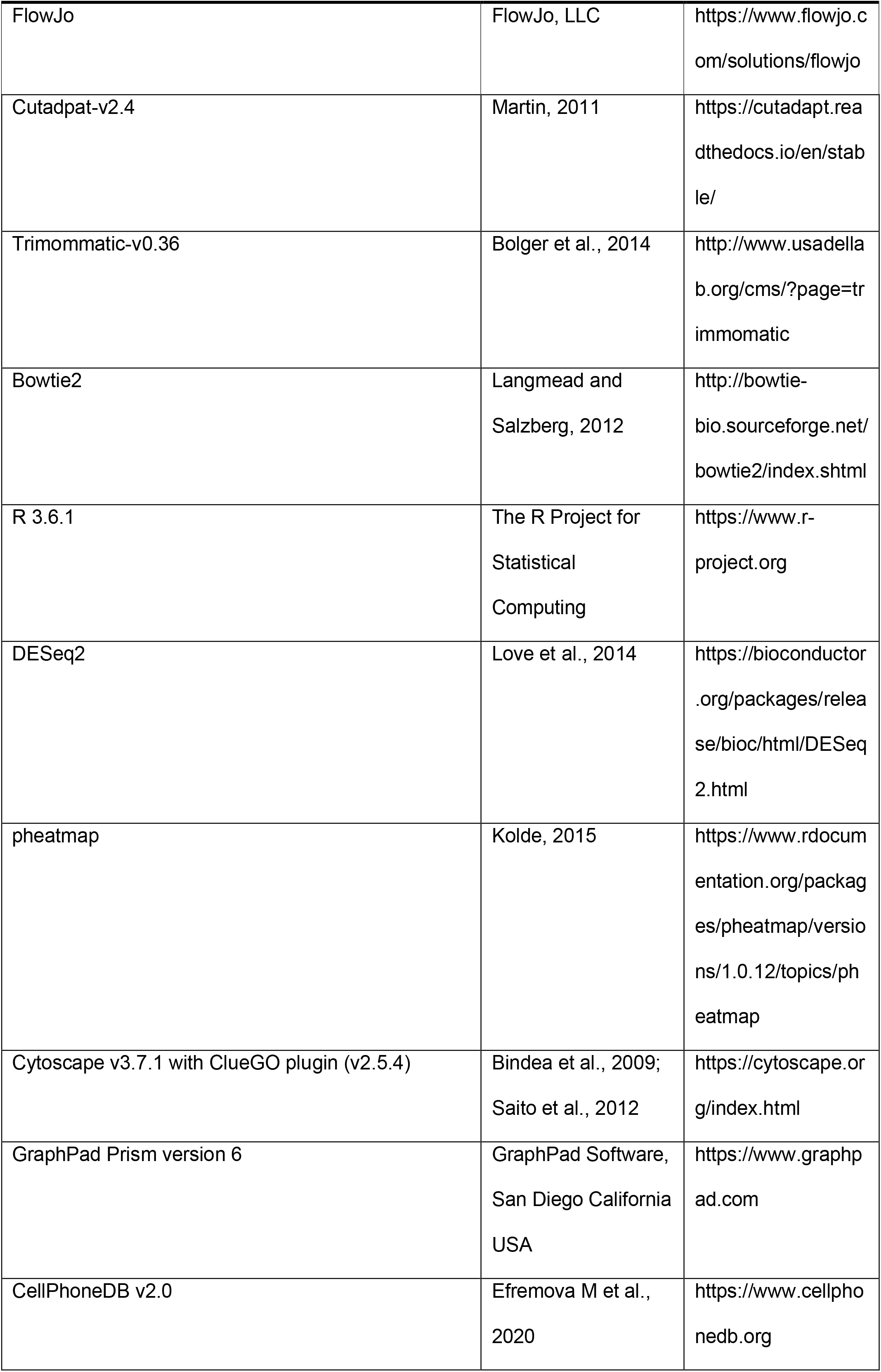
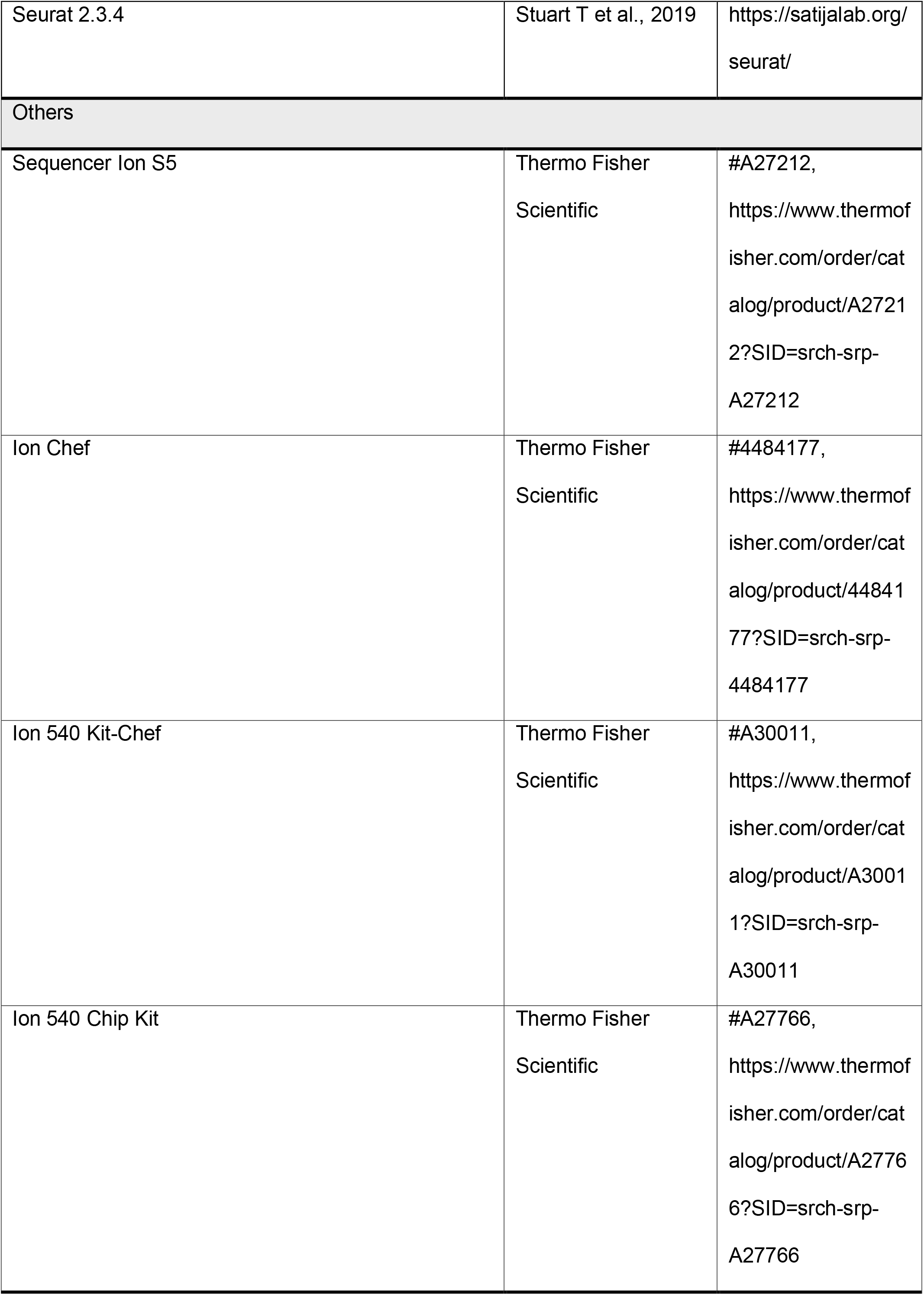

## Supplemental Information

**Figure S1.**
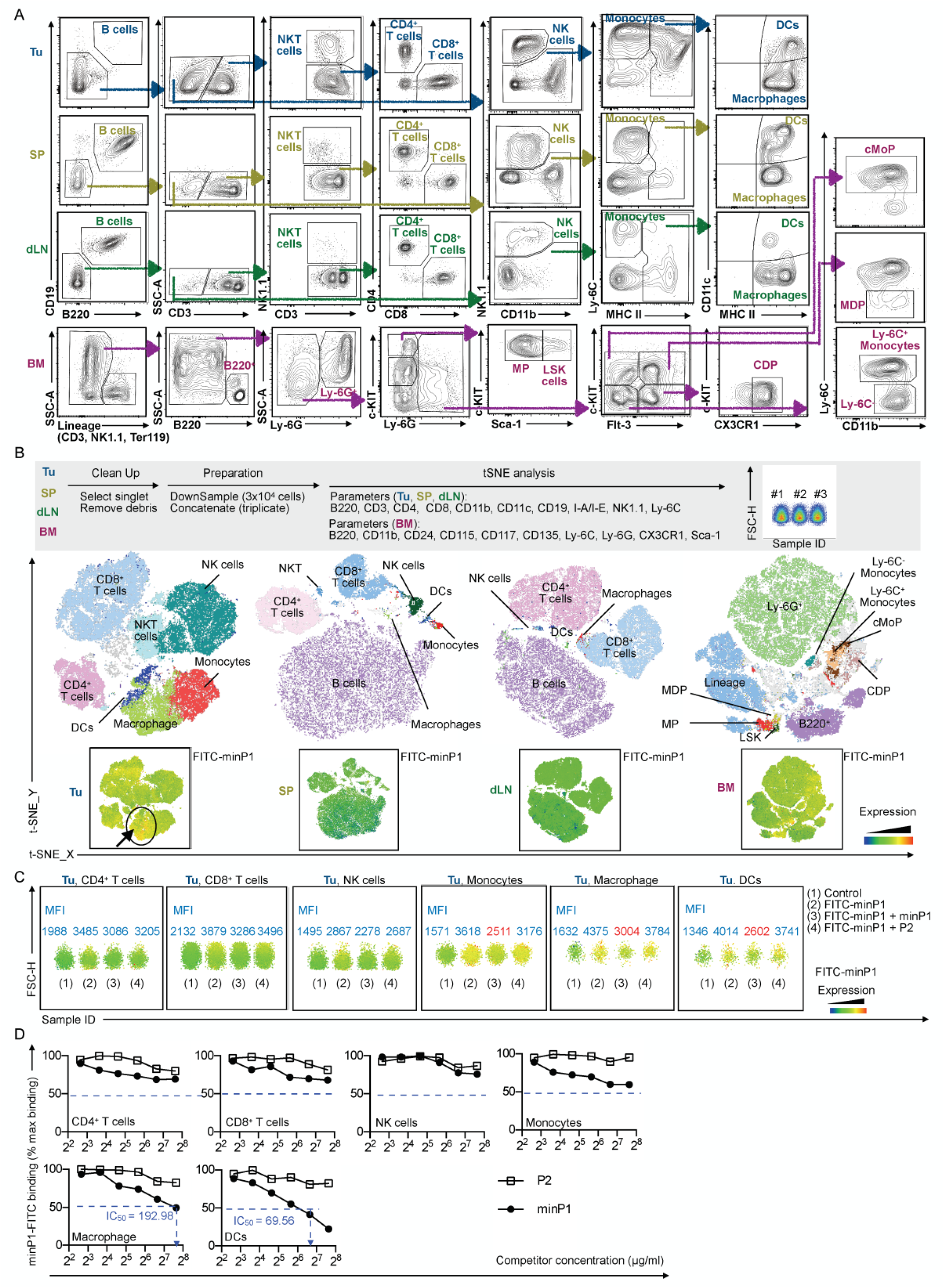
minP1 preferentially binds on intratumoral DC populations. **(A)** Flow cytometry gating of distinct immune cell populations in tumor (Tu), draining lymph node (dLN), spleen (SP), and bone marrow (BM). (**B)** The tSNE identifies distinct immune cell populations in the tumor and also displays the fluorescence intensity of FITC-minP1 in those populations. **(C)** Competitive binding assay of FITC-minP1 florescence intensity in different tumor immune cell populations. Unlabeled minP1 and the P2 (derived from the HMGN1 CHUD) act as a FITC-minP1 antagonist and noncompetitive control, respectively. **(D)** Binding affinities of FITC-minP1 for different tumor cell populations, using various doses of minP1 as a FITC-minP1 antagonist, and P2 as a noncompetitive control. Abbreviation: MFI (mean fluorescence intensity), PI (propidium iodide).

**Figure S2.**
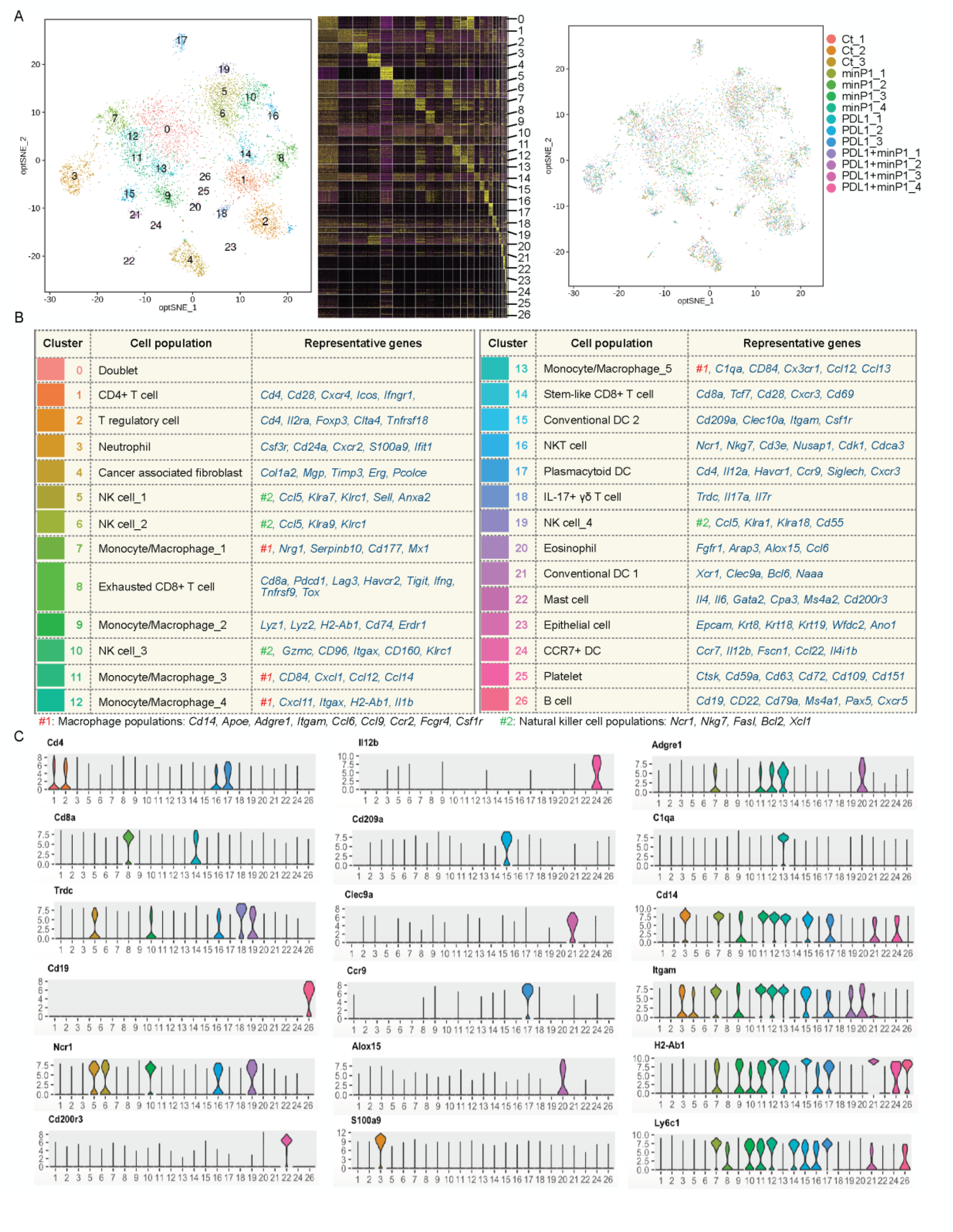
Overview of single-cell RNA sequencing results on tumors of Colon26 tumor-bearing mice after minP1/αPDL1 treatment. **(A)** A t-distributed stochastic neighbor embedding (t-SNE) projection of scRNA-seq profiling from 7613 CD45^+^ cells in tumors of Colon26 tumor-bearing mice. Cell clusters are distinct colors. **(B)** scRNA-seq sample classifications for CD45^+^ cells in different treatment groups. **(C)** Overview of selected genes in different cell clusters. **(D)** Violin plots showing the expression distribution of selected genes in each cell clusters. The y-axis represents log-normalized expression value.

**Figure S3.**
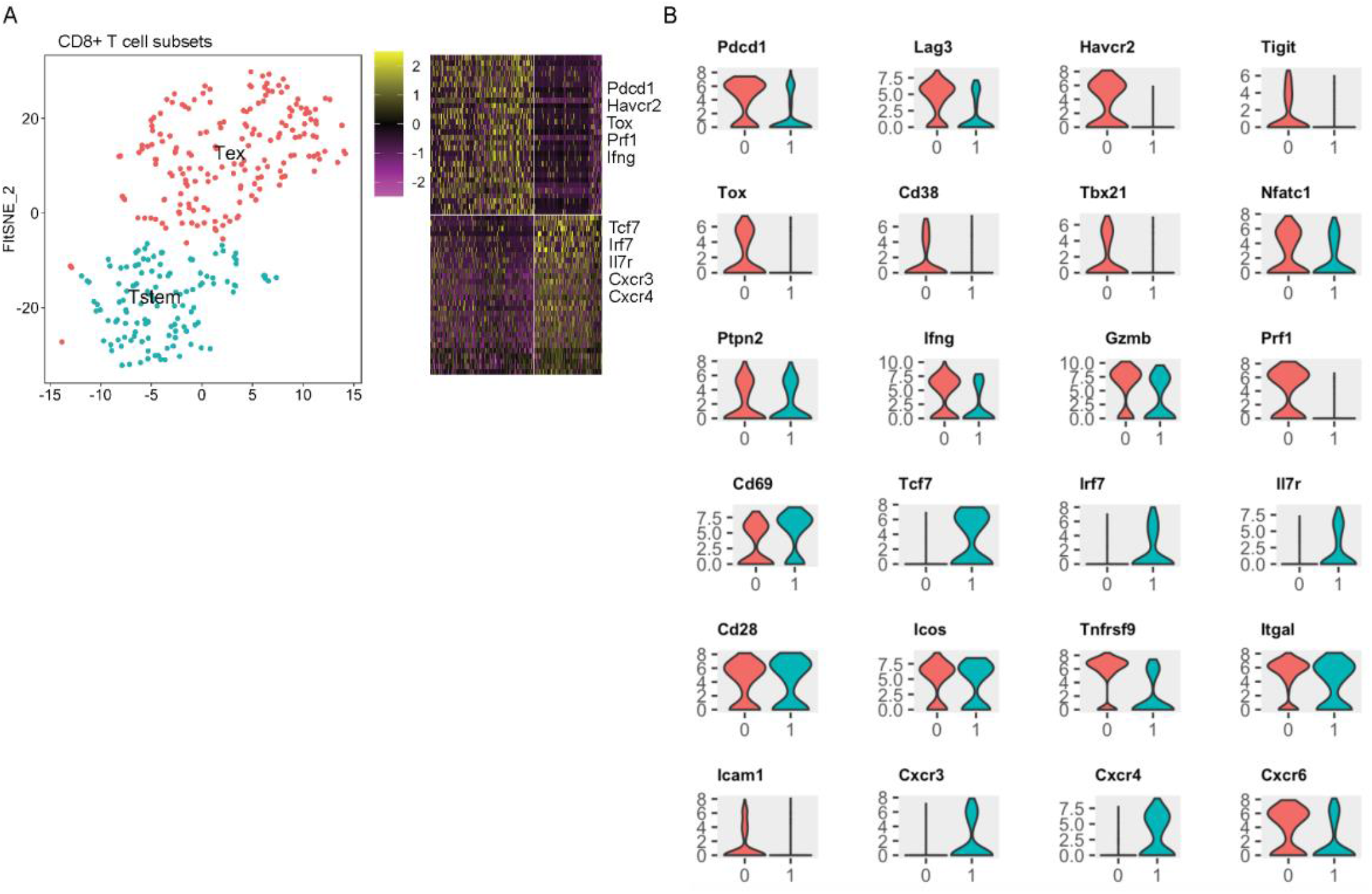
Overview of single-cell RNA sequencing results on characterization of CD8^+^ T cell subsets. **(A)** A t-distributed stochastic neighbor embedding (t-SNE) projection of scRNA-seq profiling from 303 CD8^+^ cells in tumors of Colon26 tumor-bearing mice. CD8^+^ T cell clusters are distinct colors. Terminally exhausted CD8^+^ T cells (Tex cell, *Pdcd1, Havcr2, Tox, Prf1, Ifng*) and stem-like CD8^+^ T cells (Tstem cell, *Tcf7, Irf7, Il7r, Cxcr3, Cxcr4*) were profiled. Violin plots showing the expression distribution of selected genes in different CD8^+^ cell clusters. The y-axis represents log-normalized expression value.

**Figure S4.**
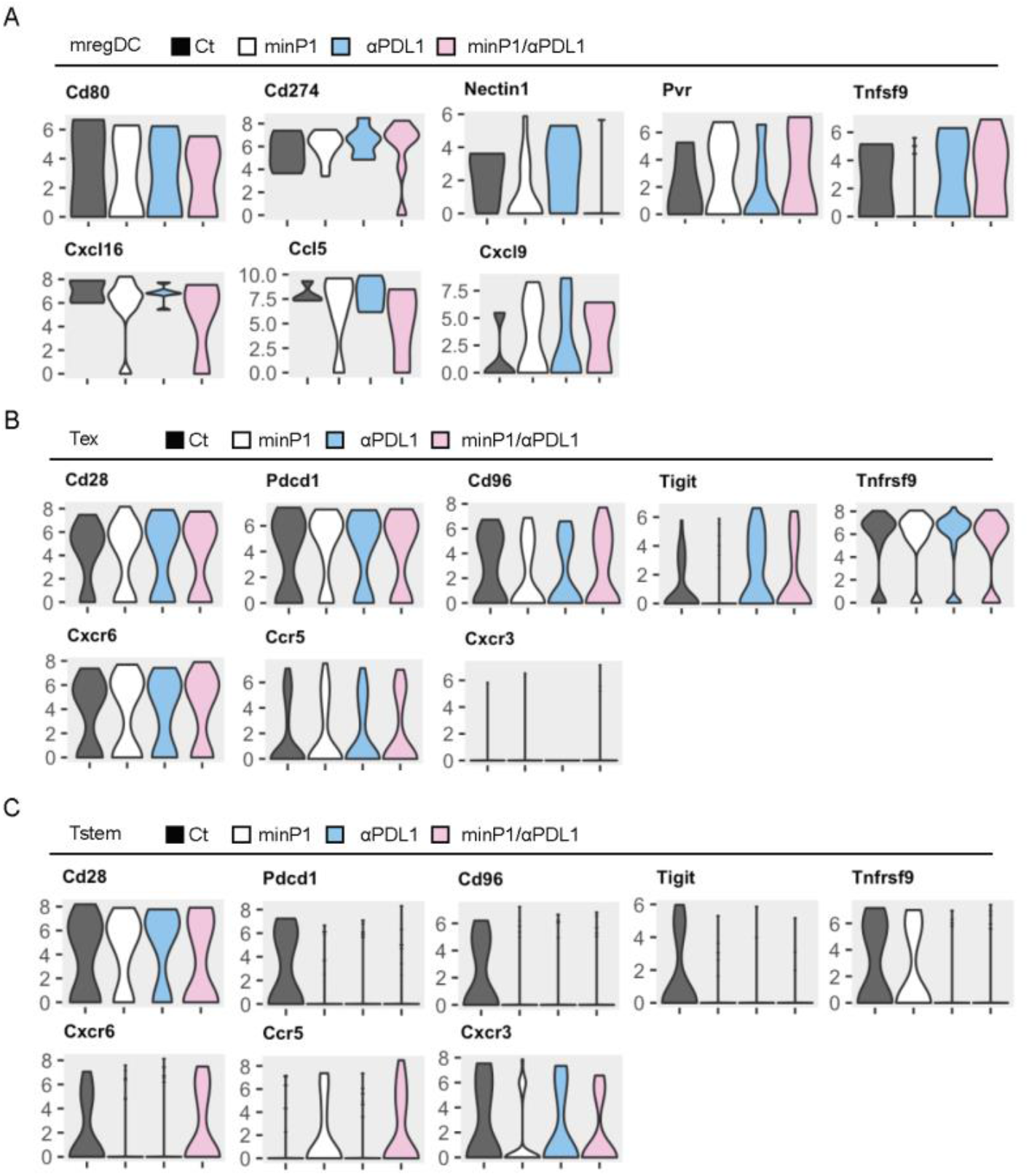
Overview of single-cell RNA sequencing results on the expression level of immunoregulatory molecules. Violin plots showing the expression of immunoregulatory molecules and their corresponding receptors or ligands on **(A)** mregDC, **(B)** Tstem cell, and Tex cell populations among different treatment groups. The y-axis represents log-normalized expression value.

